# Disulfide constrained Fabs overcome target size limitation for high-resolution single-particle cryo-EM

**DOI:** 10.1101/2024.05.10.593593

**Authors:** Jennifer E. Kung, Matthew C. Johnson, Christine C. Jao, Christopher P. Arthur, Dimitry Tegunov, Alexis Rohou, Jawahar Sudhamsu

## Abstract

High-resolution structures of proteins are critical to understanding molecular mechanisms of biological processes and in the discovery of therapeutic molecules. Cryo-EM has revolutionized structure determination of large proteins and their complexes^1^, but a vast majority of proteins that underlie human diseases are small (< 50 kDa) and usually beyond its reach due to low signal-to-noise images and difficulties in particle alignment^2^. Current strategies to overcome this problem increase the overall size of small protein targets using scaffold proteins that bind to the target, but are limited by inherent flexibility and not being bound to their targets in a rigid manner, resulting in the target being poorly resolved compared to the scaffolds^3–11^. Here we present an iteratively engineered molecular design for transforming Fabs (antibody fragments), into conformationally rigid scaffolds (Rigid-Fabs) that, when bound to small proteins (∼20 kDa), can enable high-resolution structure determination using cryo-EM. This design introduces multiple disulfide bonds at strategic locations, generates a well-folded Fab constrained into a rigid conformation and can be applied to Fabs from various species, isotypes and chimeric Fabs. We present examples of the Rigid Fab design enabling high-resolution (2.3–2.5 Å) structures of small proteins, Ang2 (26 kDa) and KRAS (21 kDa) by cryo-EM. The strategies for designing disulfide constrained Rigid Fabs in our work thus establish a general approach to overcome the target size limitation of single particle cryo-EM.

## MAIN

Technological advances in cryogenic electron microscopy (cryo-EM) have enabled the determination of high-resolution structures of large proteins and protein complexes. Solving the structures of small proteins (< 50 kDa), however, remains a major challenge, as their lack of distinctive low-frequency structural features and the low signal-to-noise ratio in cryo-EM images prevents accurate single-particle image alignment. Yet, most proteins in both prokaryotic and eukaryotic cells, including many drug targets, are smaller than 50 kDa^12^. Consequently, this powerful technique remains largely inaccessible for a vast proportion of the proteome.

Several approaches have been proposed to address this problem. Most of these seek to increase the apparent size of the small protein by designing another protein, a “structure chaperone”, that binds with high affinity to the small protein. Structure chaperones, such as designed ankyrin repeat proteins (DARPins), nanobodies, or Fabs, can be evolved using in vivo and in vitro methods to bind to specific targets. The small size (∼15 kDa) of DARPins and nanobodies, however, limits their use as structure chaperones for cryo-EM. To overcome this limitation, both nanobodies and DARPins have been fused or bound to other proteins to produce elaborate scaffolds, such as “legobodies”^13^, “pro-macrobodies”^8^, “megabodies”^14^, and self-assembling protein cages^4,6^, of sufficient size to visualize a small protein of interest. However, inherent flexibility of the resulting fusion proteins and assemblies^3,4,15^ limits the overall resolution of the small target protein, which is often poorly resolved (Extended Data Fig. 1).

**Figure 1.**
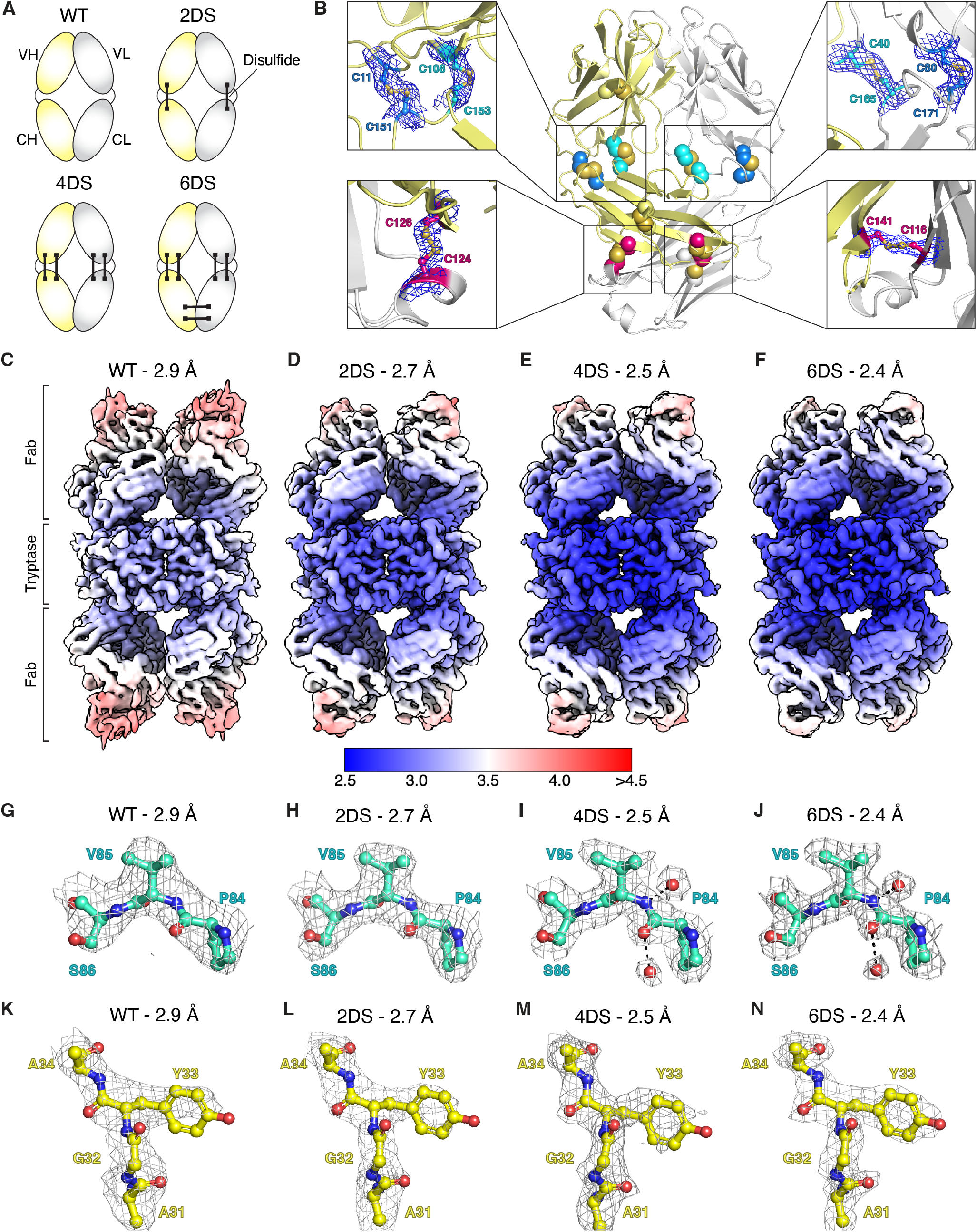
Design and characterization of rigid Fabs. **(A)** Design of Rigid Fabs. To engineer Fabs that are conformationally rigid, two (2DS) or four (4DS) intrachain disulfides were introduced in the elbow region of both the light (gray) and heavy (yellow) chains to restrict the elbow angle between the variable and constant domains. Two additional interchain disulfides were introduced in the constant domain to further reduce the flexibility of this domain, leading to a Rigid Fab design containing six engineered disulfides (6DS). **(B)** Crystal structure of E104.v1.6DS Fab. Center, cartoon representation of E104.v1.6DS (HC, yellow; LC, gray) with disulfides shown in spheres (2DS disulfides, cyan; 4DS disulfides, blue; 6DS disulfides, pink). Insets show electron density contoured at 1σ for each of the engineered disulfides. **(C-F)** maps of tryptase complexes with (C) WT, (D) 2DS, (E) 4DS, and (F) 6DS variants of the E104.v1 Fab colored by local resolution. **(G-N)** EM density for selected map regions illustrating improvement in resolution with increasingly rigid constructs of E104.v1: (G-J) tryptase P84-S86 (green sticks) and (K-N) E104.v1 HC A31-A34 (yellow sticks) from the indicated tryptase-Fab structures.

Fabs, on the other hand, possess several favorable properties that make them an attractive option as structure chaperones: (a) novel Fabs can be discovered using well-established methods to bind almost any target with high affinity; (b) at ∼50 kDa, Fabs by themselves are theoretically large enough to be resolved to high resolution via cryo-EM^16^; (c) their distinctive shape provides a recognizable feature that facilitates accurate image alignment^17^. The primary limitation of Fabs as structure chaperones for cryo-EM has been their inherent conformational flexibility^18^.

Here, we applied iterative structure-guided protein engineering to manually design >100 Fabs variants in an effort to reduce flexibility at the elbow angle, by introducing new disulfide bonds at strategic locations. These designs were first tested for expression, proper folding, and a narrow set of them resulted in well-folded Fabs and maintained affinity for the antigen. We then assessed the flexibility of the various designs by solving cryo-EM structures of relatively large antigen-Fab complexes and arrived at Fab designs that were translatable across species and chimeras, that rendered the Fab conformationally rigid. We then translated these designs to Fabs against two small targets, Ang2 (26 kDa) and KRAS (21 kDa) and demonstrated that Rigid Fabs can enable high-resolution (∼2.3–2.5 Å) structure determination of both small proteins using cryo-EM. These elegant solutions are transferable to practically any Fab against any target and could allow high resolution structure determination of any small protein.

### Design, iteration and characterization of Rigid Fabs

Crystal structures of Fabs have revealed that there is a high degree of variability in the elbow angle between the variable and constant domains (115-225°) (Extended Data Fig. 2A)^18^. This flexibility frequently causes the constant domain of the Fab to be poorly resolved in cryo-EM maps, and masks are often applied during refinement to exclude the Fab constant domain in order to achieve higher resolution at the Fab-antigen interface for large antigen-Fab complexes. Thus, only the ∼25 kDa variable domain of the Fab is fully utilized in these cases to aid in image alignment. If the Fab were conformationally rigid with a distinctive shape, however, including the constant domain would (a) increase the overall ordered size of the particle by ∼50 kDa, (b) alleviate the need for a mask to discount the constant domain, and (c) improve image alignment, leading to improved resolution of the constant domain and of the antigen-Fab complex overall. For a truly conformationally rigid Fab, the local resolution throughout the Fab would be expected to be relatively uniform.

**Figure 2.**
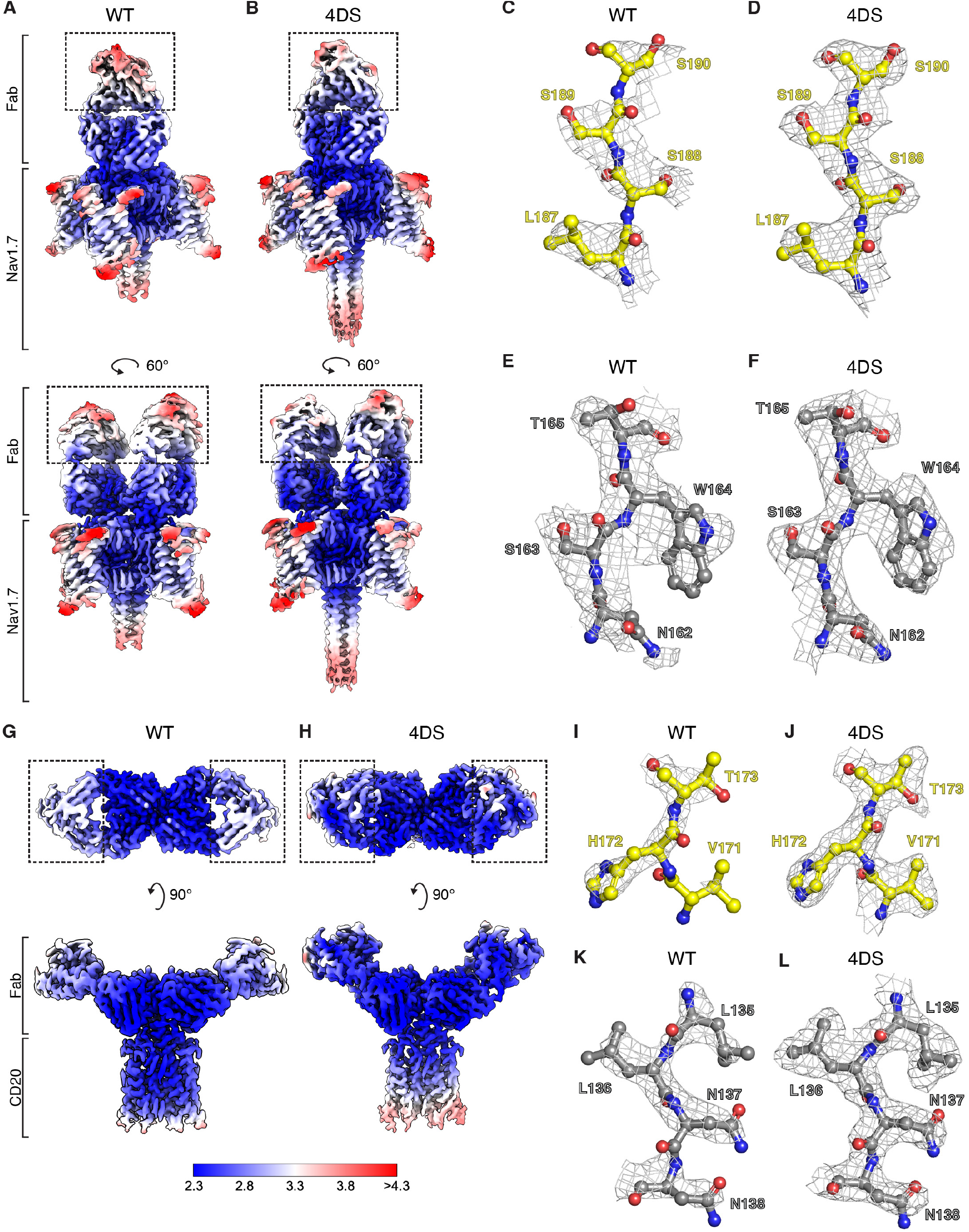
Cryo-EM structures of Nav1.7-7A9 Fab and CD20-RTX Fab complexes. **(A and B)** Composite cryo-EM maps of Nav1.7 complexes with (A) WT or (B) 4DS variants of the 7A9 Fab colored by local resolution. Dashed boxes indicate the constant domains of the Fabs. **(C-F)** EM density for selected map regions illustrating improvements in resolution in the 7A9 HC (C-D) and LC (E-F) from the indicated structures. **(G and H)** Composite cryo-EM maps of CD20 complexes with (G) WT or (H) 4DS variants of the rituximab (RTX) Fab colored by local resolution. Dashed boxes indicate the constant domains of the Fabs. **(I-L)** EM density for selected map regions illustrating improvements in resolution in the RTX HC (I-J)or LC (K-L) from the indicated structures.

Previous attempts to generate conformationally rigid Fabs utilized phage display to shorten and mutate the heavy chain (HC) elbow of the Herceptin Fab framework, and clones were selected based on their thermal stability^10^. However, use of Fabs with these modifications in cryo-EM studies revealed that they are still quite flexible^11,19–22^ (Extended Data Fig. 1F). To generate truly conformationally rigid Fabs, we sought protein engineering solutions to restrict the flexibility between the variable and constant domains that could be easily transferred between Fabs from different species, frameworks, as well as chimeric Fabs with variable and constant domains from different species.

Our analysis of structural alignments of existing crystal structures of Fabs from various species and frameworks revealed that rabbit Fabs are naturally less flexible than human or murine Fabs. (Extended Data Fig. 2A, Extended Data Table 1). This reduced flexibility is likely due to an interdomain disulfide between residues C80 and C171 (Kabat numbering) in the light chain (LC) elbow^23^. Structural analysis indicated that mutation of the corresponding residues in human Fabs, P80 and S171, to cysteine could allow for disulfide bond formation. We hypothesized that introducing additional disulfide bonds in the elbow region could result in Fabs that are conformationally even more rigid (Fig. 1A). We analyzed structures of Fabs in the Protein Data Bank (PDB) and identified additional pairs of conserved residues (E81:S168, F83:Q166, I106:S171 in the LC and L11:P151, T110:P151 in the HC, anti-Tryptase Fab in 6VVU as reference) that were in close proximity with Cβ-Cβ distance ≤ 5.5 Å and oriented such that cysteine mutations may allow formation of disulfide bonds connecting the variable and constant domains. (Extended Data Fig. 2B).

To assess the rigidity of the Fab designs, we chose the symmetric tetramer of human β-tryptase (120 kDa) and E104.v1, an anti-tryptase Fab that forms a 4:4, ∼320 kDa complex with tryptase^24^. We mutated the residue pairs mentioned above to cysteines in E104.v1 to generate Fabs that contained an elbow disulfide in their LC, HC, or both. Expression levels of the Fab variants were comparable to wild type (WT), and intact protein LC-MS indicated that each Fab had the expected additional number of disulfide bonds. Biolayer interferometry (BLI) showed that all Fab variants exhibited identical binding affinity and kinetics as E104.v1.WT and formed a 4:4 complex with tryptase (Extended Data Fig. 3).

**Figure 3.**
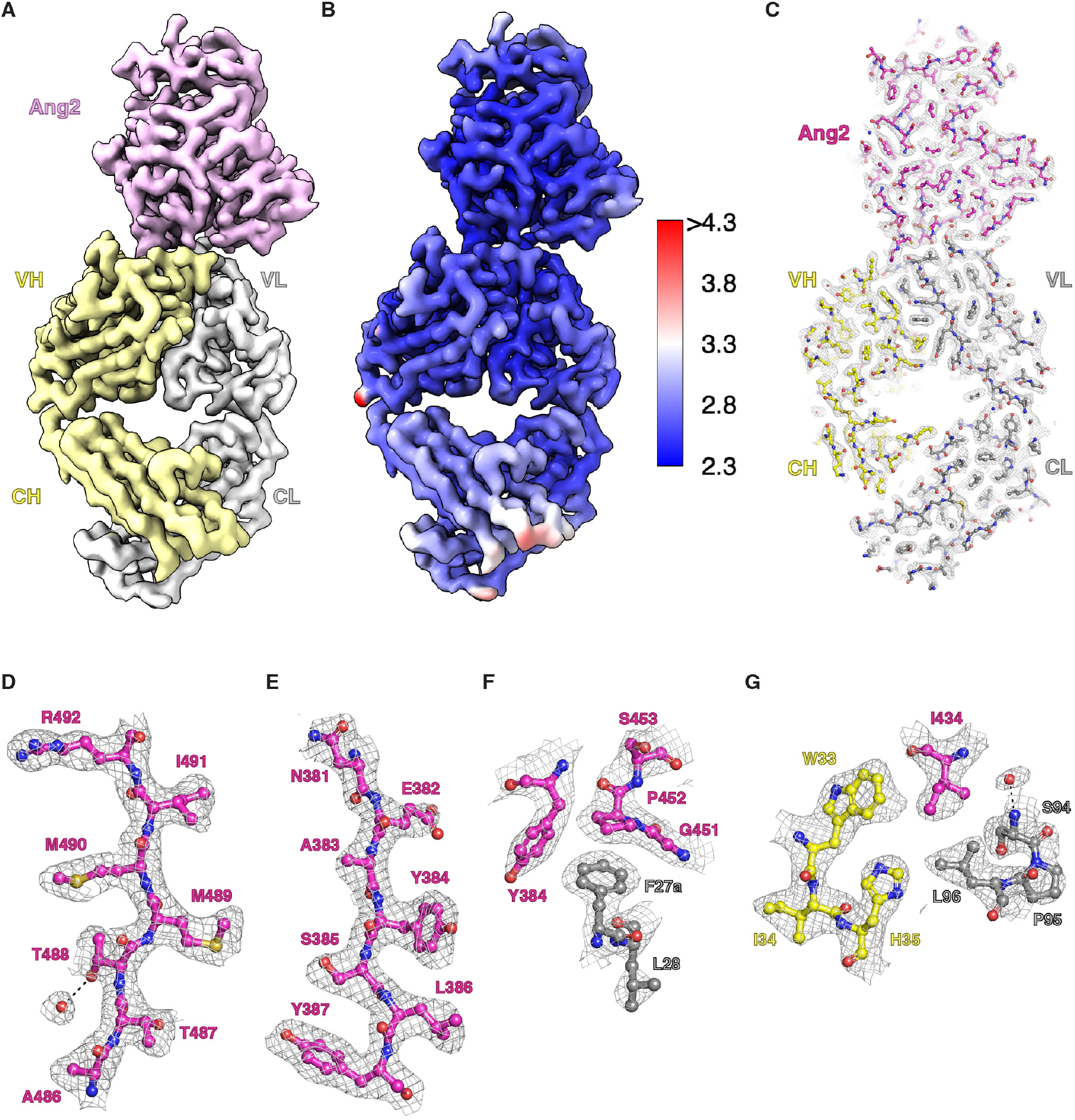
Cryo-EM structure of Ang2-5A12.6DS Fab complex. **(A)** Cryo-EM map of Ang2-5A12.6DS Fab complex at a resolution of 2.7 Å with Ang2 in pink, Fab HC in yellow, and LC in light gray. **(B)** Cryo-EM map of Ang2-5A12.6DS complex colored by local resolution. **(C-G)** EM density for the entire Ang2-5A12.6DS complex (C) and selected map regions (D-G) illustrating high resolution features, with Ang2 in magenta, RTX LC in gray, and HC in yellow.

We collected cryo-EM datasets for samples of the tryptase tetramer in complex with E104.v1.WT or with a construct containing two elbow disulfides (2DS) formed by variants L11C:P151C in the HC and P80C:S171C in the LC. For the WT dataset, we obtained a reconstruction with a resolution of 2.9 Å (Fig. 1C, Extended Data Fig. 4A-E, Extended Data Table 2). The tryptase tetramer was the best-resolved part of the structure, and the Fabs were relatively poorly resolved, especially in the constant domains, consistent with the expected flexibility of WT Fab. From the 2DS dataset, we obtained a reconstruction with an improved resolution of 2.7 Å (Fig. 1D, Extended Data Fig. 4F-J, Extended Data Table 2) and higher local resolution in tryptase, as well as the Fab variable and constant domains, compared to the WT structure, indicating that the 2DS Fab was indeed less flexible than the WT Fab (Fig. 1C-D, 1G-H, 1K-L).

**Figure 4.**
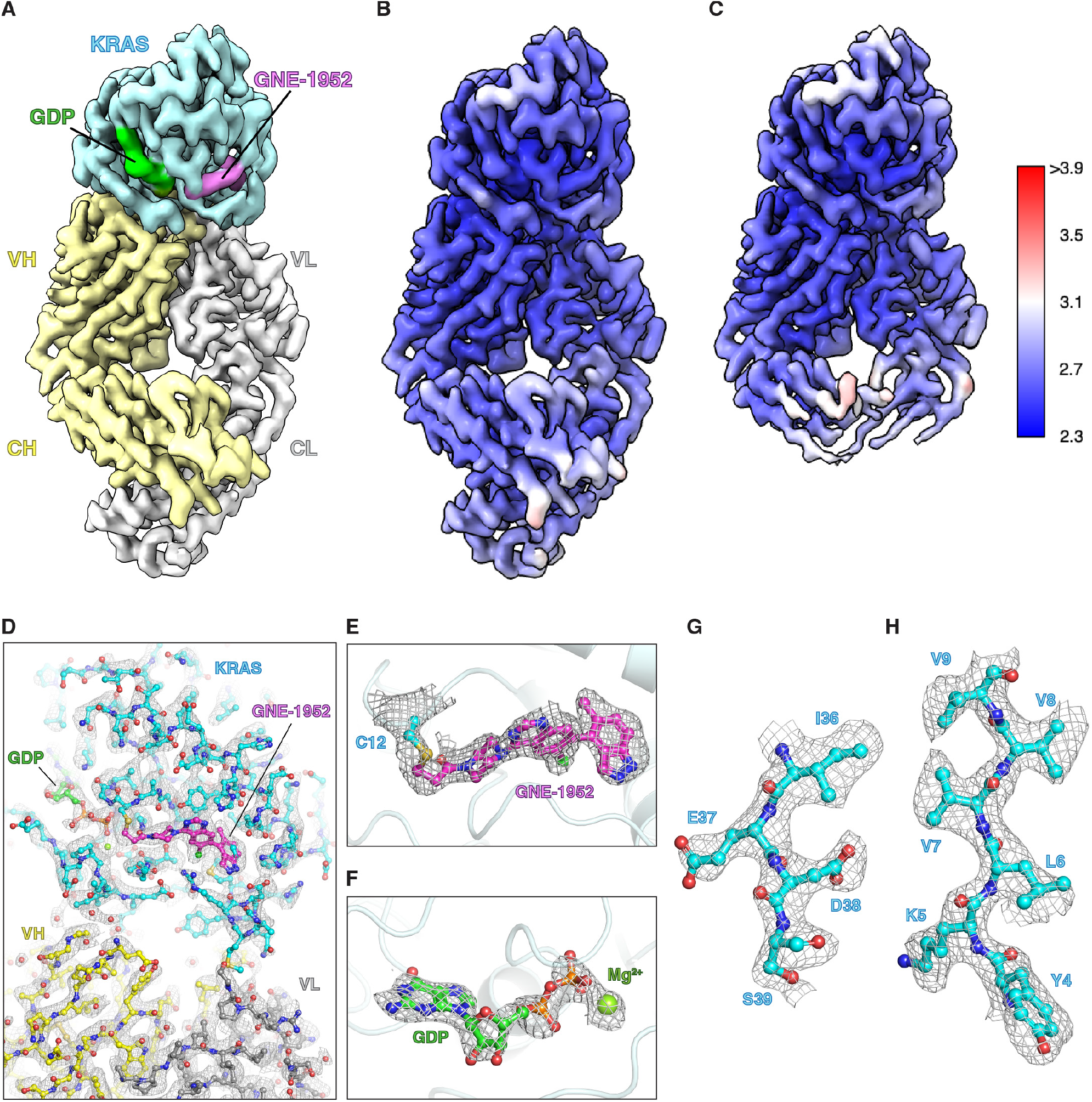
Cryo-EM structure of KRASG12C-GNE-1952-2H11.4DS Fab complex. **(A)** Cryo-EM map of KRAS^G12C^-GNE-1952-2H11.4DS Fab complex at a resolution of 2.8 Å with KRAS^G12C^ in cyan, GNE-1952 in pink, GDP in green, 2H11.6DS HC in yellow, and LC in light gray. **(B)** map of KRAS^G12C^-GNE-1952-2H11.4DS complex colored by local resolution. **(C)** Locally refined cryo-EM map of KRAS^G12C^-GNE-1952-2H11.4DS complex colored by local resolution. **(D-H)** EM density for the entire KRAS^G12C^-GNE-1952-2H11.4DS complex (D), GNE-1952 covalently bound to C12 (E), GDP (F), and selected regions of KRAS^G12C^ (G-H) illustrating high resolution features.

Though the 2DS Fab was more rigid than the WT Fab, the constant domains of the 2DS Fab were still less well resolved than the variable domains, indicating some flexibility remained in the 2DS Fab elbow. To generate Fabs that are more rigid, we identified additional sites in the Fab LC and HC elbows to introduce more disulfides to further constrain the conformation of the Fab (Fig. 1A, Extended Data Fig. 2C). The structure of the E104.v1.WT-tryptase complex (PDB code 6VVU) indicated that the mutations P40C:E165C in the LC could lead to disulfide formation. In the HC, L108 and P153 were too far apart (Cβ-Cβ distance = 6.3 Å) to allow for disulfide formation with L108C:P153C mutations. Since P153 is in a loop, we inserted a cysteine residue between E152 and P153 to lengthen the loop and position the new cysteine close enough to enable disulfide bond formation with L108C. This design, 4DS, contained a total of four intrachain disulfides in the elbow region, two in each chain (Fig. 1A). As with the 2DS Fabs, the various 4DS Fabs expressed at levels similar to E104.v1.WT, possessed all four engineered elbow disulfides as indicated by intact LC-MS analysis, and bound to tryptase with similar binding affinity and kinetics as E104.v1.WT (Extended Data Fig. 3). To further characterize these Fabs, we crystallized and solved structures of the 4DS Fab variants at a resolution of 2.0 Å (Extended Data Fig. 2E-F, Extended Data Table 3). The structures confirmed that all four elbow disulfides formed as designed. To determine whether the additional elbow disulfides in the 4DS Fab would further reduce flexibility, we collected a cryo-EM dataset for the tryptase-4DS Fab complex and obtained a 3D reconstruction with a resolution of 2.5 Å (Fig. 1E, Extended Data Fig. 4K-O, Extended Data Table 2). Strikingly, in addition to the improved overall resolution, we observed notable improvements in local resolution throughout the structure including several well-resolved water molecules (Fig. 1E, 1I, 1M), and especially in the constant domains of the Fabs, demonstrating that the new disulfides in the 4DS Fab significantly increased the rigidity of the Fab.

We observed that the C-termini of the constant domains of the 4DS Fabs still retained some flexibility relative to the rest of the Fab (Fig. 1E) and sought to engineer this region of the Fab to further decrease flexibility. We examined our 4DS Fab crystal structures and identified additional sites in the constant domain where we could introduce interchain disulfides between the LC and HC (Extended Data Fig. 2D). We designed and expressed variants of E104.v1 that contained pairs of cysteine mutations at one or more of these sites in combination with the 4DS elbow mutations. All of the 5DS/6DS Fabs bound to tryptase with similar binding affinity and kinetics as E104.v1.WT (Extended Data Fig. 3), demonstrating that the constant domain mutations also do not negatively affect the ability of the Fab to bind antigen. Furthermore, crystal structures of a 5DS (F116CLC:A141CHC) and a 6DS (F116CLC:A141CHC and Q124CLC:F126CHC) Fab further confirmed proper formation of all engineered disulfides (Fig. 1B, Extended Data Fig. 2G, and Extended Data Table 3). We collected a cryo-EM dataset for the tryptase-6DS Fab complex and obtained a 3D reconstruction with an overall resolution of 2.4 Å (Fig. 1F, Extended Data Fig. 4P-T, Extended Data Table 2). Overall, the 6DS map was very similar to the 4DS map with some improvements in local resolution, especially at the C-termini of the constant domains, consistent with the interchain disulfides in the constant domain increasing the rigidity of that domain (Fig. 1F, 1J, 1N). Comparison of ResLog plots for each of the tryptase-Fab complexes further demonstrated that each additional pair of engineered disulfides in the Fab led to marked improvements in the quality of the data, while also reducing the number of particles required to achieve higher resolutions (Extended Data Fig. 4U)^25^.

### Modularity of Rigid Fab designs

In light of our success designing rigid versions of E104.v1, a fully humanized Fab, we speculated that the same designs could be applied to rigidify Fabs derived from other species or chimeras. We selected the murine Fab 7A9 and the human-mouse chimeric Fab fragment from Rituximab (RTX), which target the membrane proteins Nav1.7 and CD20, respectively. Previous cryo-EM structures of these WT Fabs bound to Nav1.7 and CD20 revealed inherent flexibility in the Fabs^26,27^. We generated 4DS variants of both 7A9 and RTX Fabs, using cysteine mutations at the same exact positions (Kabat numbering) as those introduced in the E104.v1.4DS Fab. Intact LC/MS analysis indicated that all four elbow disulfides were formed in both 7A9.4DS and RTX .4DS Fabs. We then collected cryo-EM datasets for each antigen bound to either the WT Fab or 4DS Fab (Fig. 2, Extended Data Fig. 5). For Nav1.7, which forms a 4:2 Nav1.7:Fab complex, the overall resolution of the consensus 3D reconstructions for both the Nav1.7-7A9.WT and Nav1.7-7A9.4DS complexes was 2.6 Å (Fig. 2A-B, Extended Data Fig. 5-6, Extended Data Table 4). Local refinement with a mask around the Fabs in the 7A9.4DS complex led to greater improvements in local resolution compared to 7A9.WT, especially in the constant domain, consistent with 7A9.4DS having increased rigidity (Fig.2A-F, Extended Data Fig. 5).

For CD20, which forms a 2:2 CD20-RTX Fab complex, the overall resolution of the consensus 3D reconstructions for both the CD20-RTX.WT and CD20-RTX.4DS complexes was 2.6 Å (Fig.2G-H, Extended Data Fig. 6, Extended Data Table 4). The constant domains of the 4DS Fabs were resolved to a much higher resolution than the WT Fabs, confirming that the 4DS mutations did indeed render the Fab significantly more rigid (Fig. 2G-L). This increased rigidity of RTX.4DS appeared to lead to lower resolution of the portions of the CD20 transmembrane helices distal to the Fab binding site, compared to the map obtained for the RTX.WT complex (Extended Data Fig. 6). We suspect that this effect was in part due to the nature of the CD20-RTX complex, in which two Fabs bind to one side of the CD20 dimer and to each other. The rigidity and greater mass of the 4DS Fabs, combined with the relative intrinsic flexibility of CD20 and added alignment noise from the detergent micelle surrounding it, likely caused the Fabs to become very well aligned at the cost of alignment accuracy and resolution in the regions of CD20 distal from the Fabs during refinement. In the case of the WT complex, the flexibility of the Fabs caused the constant domains to have less weight in determining particle alignments, allowing for improved alignments for CD20 compared to the 4DS Fab complex. Iterative local refinements using masks to exclude the 4DS Fab constant domains, followed by exclusion of the entire Fab, led to significant improvements in the resolution of the intracellular side of the CD20 helices (Extended Data Fig. 6L). The consensus map and focused maps were combined to generate a composite map that had improved resolution at the Fab-CD20 interface, as well as throughout CD20. The same procedure was applied to the RTX.WT dataset, leading to a final composite map with similar overall resolution to the RTX.4DS composite map (Fig. 2G-H, Extended Data Fig. 6). Thus, even though the rigidity of the 4DS Fab initially led to lower resolution of the antigen, local refinements were able to ameliorate this effect. While this effect was not ideal for solving a structure of this particular antigen, our observation that the RTX.4DS Fab could so strongly drive particle alignments underscored the marked increase in rigidity of 4DS Fab variants relative to WT Fabs.

Overall, the increased rigidity observed for both the 4DS variants of the 7A9 and RTX Fabs demonstrated the modularity of the 4DS Rigid Fab design. The same set of mutations introduced in the humanized E104.v1.4DS Fab successfully resulted in conformationally rigid Fabs against other antigens, even for murine or chimeric Fabs like 7A9 and RTX, respectively.

### Rigid Fabs enable high-resolution structures of small proteins

To date, there are no cryo-EM structures of Fabs in complex with monomeric antigens smaller than 50 kDa in the EMDB with resolutions better than ∼3.0 Å, and only 7 unique monomeric antigen-Fab complexes with resolutions better than ∼3.5 Å, underscoring the difficulty of using flexible WT Fabs to solve high-resolution structures of small antigens. In addition, there are no structures currently in the EMDB of monomeric antigens smaller than 50 kDa bound to any structure chaperone that have resolutions better than ∼3.0 Å. We next sought to determine whether Rigid Fabs, given their moderate size, were sufficient to enable high resolution structures of small antigens. To test this, we selected the cytokine Ang2 (26 kDa), an extracellular protein and the GTPase KRAS (21 kDa), an intracellular protein, both of which are monomeric, as targets and generated a 6DS variant of the anti-Ang2 Fab 5A12 and a 4DS variant of the anti-KRAS Fab 2H11 based on our designs for the 4DS and 6DS variants of the anti-tryptase Fab E104.v1^28,29^.

### High-resolution cryo-EM structure of a 26 kDa cytokine

To evaluate if we could use Rigid Fabs to solve a high resolution cryo-EM structure of Ang2, we first generated a 6DS variant of the 5A12 Fab by introducing cysteine mutations at the same positions as in the E104.v1.6DS Fab and assembled a 74 kDa Ang2-5A12.6DS Fab complex. To further increase the mass of the sample and improve our chances at obtaining a high resolution structure, we also generated a fusion of the Protein A D domain to Protein G (ProA-ProG) and formed a ternary complex with 5A12.6DS bound to Ang2 (Extended Data Fig. 7A). We then collected a cryo-EM dataset for this complex, but 2D class averages revealed that the vast majority of the particles contained only Ang2 and the 6DS Fab, suggesting that the affinity of ProA-ProG was not high enough to remain bound at the low sample concentrations used during grid freezing (Extended Data Fig. 7B). Despite the small size (74 kDa) of the Ang2-5A12.6DS complex, secondary structure in both Ang2 and the Fab were surprisingly clearly resolved in the 2D class averages. Through iterative rounds of heterogeneous refinement, particles containing ProA-ProG were removed, leaving a particle stack containing only the Ang2-5A12.6DS complex. These particles were then used to generate a 3D reconstruction with an overall resolution of 2.7 Å, with most regions of Ang2 reaching higher resolutions, up to 2.3 Å (Fig. 3B, Extended Data Fig. 7C-E, Extended Data Table 5). The local resolution is relatively uniform (∼2.4-2.8 Å) throughout Ang2 and the Fab, consistent with 5A12.6DS being conformationally rigid (Fig. 3B). The overall structure of Ang2 is nearly identical to that observed in the 2.3 Å crystal structure of Ang2 and 5A12.WT (RMSD = 0.36 Å)^28^. The resolution of the map enabled unambiguous placement of side chains in Ang2, the Fab, and the Ang2-Fab interface, and of several water molecules (Fig. 3C-G), highlighting that using a Rigid Fab design enabled high-resolution structure determination of the 26 kDa Ang2 by cryo-EM.

### High-resolution cryo-EM structure of a small molecule bound to 21 kDa KRAS

Next, we reasoned that Rigid Fabs could be used to accelerate structure-based drug design efforts by enabling high-resolution cryo-EM structure determination of small molecule drug targets, many of which are < 50 kDa and sometimes can be intractable for structure determination by crystallography or cryo-EM. We chose the 21 kDa GTPase KRAS, for which a Fab has been reported29. Unlike the Fabs described so far in the current work, the anti-KRAS Fab 2H11 contains a lambda LC rather than a kappa LC. Lambda LCs are slightly longer, which leads to a wider range of possible elbow angles^18^. Analysis of published crystal structures of 2H11 Fab-KRAS^G12C^ complexes indicated that this Fab is indeed highly flexible29 and that the LC cysteine mutations used in our previous 2DS and 4DS designs would likely not be compatible with elbow disulfide formation in this Fab. To form the disulfide corresponding to the LC disulfide introduced in the 2DS design, we inserted a cysteine between N170 and N171 rather than a N171C mutation, as N171 was likely too far from S80 for disulfide formation (Extended Data Fig. 8A). For the second LC disulfide, we used the mutation K166C instead of Q167C since K166 was closer to P40 (Extended Data Fig. 8A). Using these mutations, we generated constructs for 4DS and 6DS variants of 2H11. While the 2H11.6DS Fab did not express, we were able to express and purify 2H11.4DS, and LC/MS indicated that all four elbow disulfides had formed. We proceeded to use 2H11.4DS to form a complex with KRAS^G12C^ conjugated to the covalent inhibitor GNE-1952 and collected a cryo-EM dataset (Extended Data Fig. 8). This led to a 3D reconstruction with an overall resolution of 2.8 Å for the whole complex, with regions of KRAS and the Fab variable domain having local resolutions of ∼2.5 Å (Fig. 4A-C, Extended Data Fig. 8C-H, Supplementary Table 5). The local resolution of the Fab constant domain was marginally lower than in the variable domain, unlike 5A12.6DS, consistent with 4DS Fabs being slightly more flexible than 6DS Fabs. Local refinement with a mask to exclude the constant domain led to an overall resolution of 2.7 Å with improvements in local resolution in KRAS, the Fab variable domain, and the ligand-binding sites, with several regions reaching ∼2.3 Å resolution (Fig. 4C). The overall structure of KRAS, GDP, and GNE-1952 in our work is nearly identical to the published crystal structure (RMSD = 0.67 Å)^29^. Importantly, the high resolution features at 2.3 Å in the ligand-binding sites for GDP and the inhibitor GNE-1952 enabled unambiguous placement of both ligands, as well as the covalent linkage between C12 and GNE-1952 (Fig. 4D-F). Side-chains throughout the KRAS^G12C^ portion of the structure were also very well-resolved (Fig. 4D, and Fig. 4G-H).

## Discussion

Computational methods for structure determination such as AlphaFold^30,31^, allow valuable modeling of the overall structures of proteins and protein complexes. Such modeling is now routinely being used to generate useful hypotheses to advance our understanding of interactions of proteins with other molecules and to expedite structure determination by providing initial models as starting points for experimental maps from either crystallography or cryo-EM. However, these models do not replace experimental structure determination^32^. The need for accurate, experimental structure determination of proteins and protein complexes to high resolution continues to exist. Small proteins (< 50 kDa) encompass a vast majority of all known proteins, especially in the context of drug targets in humans and pathogens. While several of these proteins are amenable to high-resolution structure determination by x-ray crystallography, many others have had limited success due to low expression, low solubility, or lack of crystallizability. Cryo-EM circumvents these challenges, because it requires low protein amounts and concentrations, yet remains challenging when working with small proteins, which are plagued by low signal-to-noise ratio and lack distinctive features for particle alignment. In this study, we demonstrate that Rigid Fabs increase the effective size of the target protein in a rigid manner and improve particle alignment and pose assignment. This results in high-resolution structure determination of proteins as small as ∼21 kDa, revealing features, such as water molecules and unambiguous placement of specific conformations of protein side chains, atoms, and small ligands, like inhibitors and co-factors. The modularity of our Rigid Fab designs allows their straightforward application to nearly any existing Fab or newly discovered Fab independent of its source and against any target, making this powerful tool accessible to all researchers. This modularity would especially be impactful in a drug discovery setting allowing: (a) faster structure determination of antigen-fab complex structures and epitope mapping for improved antibody discovery and optimization in early stages of immunization campaigns; and (b) iterative structure based drug design for small molecule drug targets that cannot be enabled using x-ray crystallography.

The modularity of the Rigid Fab designs is made possible by the highly conserved fold and architecture of Fabs across different species, precluding the need for a pre-existing structure of the Fab of interest in order to design a rigid version of the Fab. Sequence alignments with the Rigid Fabs presented in this study (Extended Data Fig. 9A-B) should be sufficient in most cases to identify sites to introduce the cysteine mutations needed to form disulfides to rigidify other Fabs. Furthermore, our Rigid Fab designs have resulted in Fabs that adopt a specific conformation (Extended Data Fig. 9C). The inherent flexibility of Fabs likely makes it possible for most Fabs to sample this conformation during expression and folding, enabling disulfide bond formation using the set of elbow and constant domain mutations from our designs. Our data suggests that the introduction of the described four disulfide bonds (4DS) in the Fab elbow region should suffice for high-resolution (∼2.5 Å) structure determination of a small protein, but that introduction of six disulfide bonds (6DS) can further improve the resolution.

Rigid Fabs are likely to be most transformative in the study of small (<50 kDa) protein targets. For larger targets (e.g. ∼130 kDa Nav1.7 tetramer or ∼45 kDa CD20 dimer), which likely feature more intrinsic flexibility than the Fab itself, we anticipate that Rigid Fab technology can also facilitate high-resolution structure determination at the Fab-antigen interface but may still need to be coupled with image processing strategies, such as focused refinements or flexibility analysis in order to achieve high resolutions throughout a large, flexible target. Alternatively, or as a complement to such an approach, one may deploy multiple Rigid Fabs against epitopes present on opposite “sides” of the target, in which case, the Fabs would aid not only in particle alignment, but also in 3D classification or flexibility analysis. None of these approaches should be needed, however, when the target proteins are suitably small and rigid. By enabling structure determination of targets previously intractable by cryo-EM or crystallography, most notably those with molecular weights < 50 kDa, Rigid Fab technology can now accelerate basic research into molecular mechanisms of action of proteins involved in pathways of interest, as well as speed up structure-guided drug discovery and optimization.

## ACKNOWLEDGEMENTS

The authors would like to thank the Antibody Engineering, Protein Chemistry, Biomolecular Resources Departments and the cryo-EM team at Genentech for their support in generating constructs, protein purification, and data collection; and Stanford Synchrotron Radiation Lightsource (SSRL) and Advanced Light Source (ALS) for access to the synchrotrons. Use of the Stanford Synchrotron Radiation Lightsource, SLAC National Accelerator Laboratory, is supported by the US Department of Energy, Office of Science, Office of Basic Energy Sciences under Contract No. DE-AC02-76SF00515. The SSRL Structural Molecular Biology Program is supported by the DOE Office of Biological and Environmental Research, and by the National Institutes of Health, National Institute of General Medical Sciences (P41GM103393). The contents of this publication are solely the responsibility of the authors and do not necessarily represent the official views of NIGMS or NIH. Beamline 5.0.2 of the Advanced Light Source, a DOE Office of Science User Facility under Contract No. DE-AC02-05CH11231, is supported in part by the ALS-ENABLE program funded by the National Institutes of Health, National Institute of General Medical Sciences, grant P30 GM124169-01.

## AUTHOR CONTRIBUTIONS

JEK and JS conceived and designed protein engineering strategies and experiments to generate and characterize Rigid Fabs. JEK performed recombinant protein production, binding assays, crystallization, cryo-EM sample and grid preparation, and solved structures. MCJ and CPA performed grid preparation and optimization, cryo-EM data collection, and assisted in data processing. CCJ prepared protein samples for CD20 and Nav1.7 complexes. DT and AR advised on experimental designs and cryo-EM data processing strategies. JEK and JS analyzed all data and wrote the manuscript with input from all authors.

## COMPETING FINANCIAL INTERESTS

All authors are employees of Genentech, Inc.

## DATA AVAILABILITY

Coordinates and related data for the crystal structures of the E104.v1.4DS.S112F, E104.v1.4DS.A114F, E104.v1.5DS, and E104.v1.6DS have been deposited in the PDB with the following accession codes, respectively: 8VEG, 8VGE, 8VGF, and 8VGG. Coordinates and cryo-EM maps for the structures of tryptase in complex with E104.v1.WT, E104.v1.2DS, E104.v1.4DS, and E104.v1.6DS have been deposited in the PDB and EMDB with the following accession codes, respectively: PDB 8VGH/EMD-43200, PDB 8VGI/EMD-43201, PDB 8VGJ/EMD-43202, and PDB 8VGK/EMD-43203. Coordinates and cryo-EM maps for the structure of Nav1.7 in complex with wild type 7A9 Fab have been deposited in the PDB and EMDB with the following accession codes: PDB 8VGL (model), EMD-43204 (composite map), EMD-43205 (consensus map), EMD-43206 (focused refinement of 7A9 Fab), and EMD-43207 (focused refinement of Nav1.7). Coordinates and cryo-EM maps for the structure of Nav1.7 in complex with 7A9.4DS Fab have been deposited in the PDB and EMDB with the following accession codes: PDB 8VGM (model), EMD-43208 (composite map), EMD-43209 (consensus map), EMD-43210 (focused refinement of 7A9 Fab), and EMD-43211 (focused refinement of Nav1.7). Coordinates and cryo-EM maps for the structure of CD20 in complex with wild type Rituximab Fab have been deposited in the PDB and EMDB with the following accession codes: PDB 8VGN (model), EMD-43212 (composite map), EMD-43213 (consensus map), EMD-43214 (focused refinement of Fab variable domain and CD20), and EMD-43215 (focused refinement of CD20). Coordinates and cryo-EM maps for the structure of CD20 in complex with Rituximab.4DS Fab have been deposited in the PDB and EMDB with the following accession codes: PDB 8VGO (model), EMD-43216 (composite map), EMD-43217 (consensus map), EMD-43218 (focused refinement of Fab variable domain and CD20), and EMD-43219 (focused refinement of CD20). Coordinates and cryo-EM maps for the structure of Ang2 in complex with 5A12.6DS have been deposited in the PDB and EMDB with the following accession codes: PDB 8VGP, EMD-43220. Coordinates and cryo-EM maps for the structure of KRAS^G12C^ in complex with 2H11.4DS have been deposited in the PDB and EMDB with the following accession codes: PDB 8VGQ, EMD-43221.

## Notes

### Competing Interest Statement

The authors have declared no competing interest.

