## Supplementary Data and Methods for "Disulfide constrained Fabs overcome target size limitation for high-resolution single-particle cryo-EM"

Recombinant protein expression and purification

*Wild type and Rigid Fabs*

His-tagged heavy chain and untagged light chain expression constructs were generated by gene synthesis (Genscript). Fabs were expressed by transient transfection in CHO cells, and the His-tagged Fabs were purified by nickel affinity chromatography (Histrap Excel, Cytiva) followed by size exclusion chromatography (SEC) on a Superdex 200 column (Cytiva) equilibrated in 20 mM histidine acetate pH 5.5, 150 mM NaCl. The presence of the engineered disulfides in the Rigid Fab constructs was confirmed by intact protein LC/MS.

*Tryptase*

Constructs for human β-tryptase encoding residues I31-P275 with an N-terminal His tag followed by an enterokinase (EK) cleavage site were expressed and purified from *Trichoplusia ni* insect cells as previously described[1]. Cultures were harvested 48 hours post-infection. The supernatant media was filtered through a 0.22 μm filter, and the His-tagged zymogen tryptase was purified by nickel affinity chromatography (Ni-NTA Superflow, Qiagen) followed by SEC on a Superdex 200 column (Cytiva) equilibrated in SEC buffer (10 mM MOPS pH 6.8, 2 M NaCl). Peak fractions were pooled and concentrated to 2 mg/mL prior to overnight cleavage with 0.1 mg/mL EK (New England Biolabs) at room temperature in 10 mM MOPS pH 6.8, 0.2 M NaCl, 0.5 mg/mL heparin (H3393, Sigma), which results in activation and tetramerization of tryptase. Tetrameric tryptase was purified on a Superdex 200 column equilibrated in SEC buffer.

Biotinylated tryptase was generated using a construct with a C-terminal Avi tag and coexpression with BirA to enable *in vivo* biotinylation. The biotinylated tryptase zymogen was purified via nickel affinity chromatography followed by SEC as described above. Addition of a single biotin was confirmed by intact protein LC/MS.

*Nav1.7*

Details of construct design for insect cell expression of Flag-tagged chimeric Nav1.7-NavAb were described previously[2]. Expression was performed in *T. ni* insect cells for 48 hrs. Two liters of insect cell paste were resuspended in 60 mL Cytobuster (#71009-4, EMD Millipore) supplemented with 1 μg/mL benzonase and 1x protease inhibitor (Roche). The mixture was incubated at 22°C for 5 minutes and transferred to 50mL conical tubes. 1% GDN (w/v) was added to solubilize samples. Samples were incubated with anti-Flag magnetic beads (Genscript, L00835) at 4°C for 2 hours with rotary mixing. To wash unbound proteins from the magnetic beads, a magnetic rack was used (L00723, Genscript). Beads were washed 4x with 10CV Wash Buffer containing 0.042% GDN. Proteins were eluted with Elution Buffer containing 0.042% GDN and 150ug/mL Flag peptide. Fractions were pooled and separated on a Superose 6 10/300 column (Cytiva) equilibrated in Gel Filtration Buffer (10 mM Tris pH 8.0, 100 mM NaCl, 0.042% GDN).

*CD20*

Details of construct design for insect cell expression of His-tagged CD20 were described previously[3]. Expression was performed in *T. ni* insect cells for 48 hours. Two liters of insect cell paste was resuspended in 60 mL cell resuspension buffer (25mM Tris pH 7.5, 300 mM NaCl, 10% glycerol) supplemented with 1 µg/mL benzonase and 1x protease inhibitor (Roche). The mixture was incubated at 22°C for 5 minutes and transferred to 50mL conical tubes.
1% GDN/0.1% CHS (w/v) was added to solubilize samples, and as well as washed nickel magnetic beads. Samples were incubated with Ni-charged magnetic beads (L00295, Genscript) at 4°C for 2 hours with rotary mixing. To wash unbound proteins from the magnetic beads, a magnetic rack was used (L00723, Genscript). Beads were washed 4x with 10CV Wash Buffer containing 0.02% GDN/0.002% CHS. Proteins were eluted with elution buffer containing 300mM imidazole. Fractions were pooled and separated on a Superose 6 10/300 column (Cytiva) equilibrated in gel filtration buffer (25mM Tris pH 7.5, 150mM NaCl, 0.02% GDN/0.002% CHS).

*Ang2*

C-terminally His-tagged Ang2 (residues E277-F496) was expressed and purified from *T. ni* insect cells as described previously[4]. Cultures were harvested 48 hours post-infection. The supernatant media was filtered through a 0.22 µm filter, and the protein was purified by nickel affinity chromatography (HisTrap Excel, Cytiva) followed by SEC on a Superdex 200 column (Cytiva) equilibrated in 20 mM Tris pH 7.5, 150 mM NaCl.

*ProA-ProG*

To generate an N-terminally His-tagged construct for bacterial expression of a Protein A-Protein G (ProA-ProG) fusion protein, residues F100-K153 from Protein A and residues T368-G430 from Protein G were linked with a 3xGS linker. The ProA-ProG fusion protein was expressed in BL21(DE3) cells in TB autoinduction media for 48 hours at 17°C. Cell pellets were resuspended in lysis buffer (50 mM Tris pH 8.0, 500 mM NaCl, 10% glycerol) supplemented with 20 mM imidazole, 1 EDTA-free protease inhibitor tablet, 1 µg/mL benzonase, lysis detergents (0.3 % Sb3-14 and 0.03% C7BzO), and 20 mg lysozyme. The solution was homogenized and incubated with 2 ml per 1 L pellet of Ni-charged MagBeads (L00295, Genscript) for 30 minutes at room temperature. To wash unbound proteins from the magnetic beads, a magnetic rack was used (L00723, Genscript). Beads were washed 4x with 10CV lysis buffer supplemented with 20 mM imidazole. The bound protein was eluted with lysis buffer supplemented with 300 mM imidazole. The protein was then concentrated and purified via SEC on a Superdex 75 column (Cytiva) equilibrated in 20 mM Tris pH 7.5, 150 mM NaCl.

*$KRAS^{G12C}$*

A "Cys-light" construct of $KRAS^{G12C}$ (residues M1-K169) was generated for *E. coli* expression by mutating all Cys residues except C12 to Ser. The protein was expressed in BL21(DE3) cells induced with 0.5 mM IPTG overnight at 16°C. Cells were then harvested, resuspended in lysis buffer (50 mM HEPES pH 7.0, 300 mM NaCl, 5% glycerol, 5 mM $MgCl_2$, 10 uM GDP, 1 mM

TCEP) supplemented with 1 mM PMSF, 1 µg/mL benzonase, and 1x protease inhibitor (Roche), and lysed using a microfluidizer. The clarified supernatant was passed over a NiNTA agarose column (Qiagen), and the protein was eluted with lysis buffer supplemented with 300 mM imidazole. The eluted protein was dialyzed into dialysis buffer (50 mM HEPES pH 7.0, 300 mM NaCl, 5 mM $MgCl_2$, 10% glycerol, 1 mM TCEP, 10 µM GDP) and incubated overnight with TEV protease to cleave the His tag. The sample was passed again over a Ni-NTA column to remove uncleaved protein. The flowthrough was concentrated and purified via SEC on a Superdex 75 column (Cytiva) equilibrated in SEC buffer (50 mM HEPES pH 7.0, 100 mM NaCl, 1 mM $MgCl_2$, 1 mM TCEP, 10 µM GDP).

Biolayer Interferometry

All binding assays were performed in 20 mM Tris pH 7.5, 150 mM NaCl, 0.1% BSA, 0.01% Tween20. Biotinylated tryptase was captured on streptavidin SA biosensors (Sartorius). Assays were performed in triplicate on an OctetRED384 (Fortebio). Sensorgrams were normalized to a reference well containing only buffer. Equilibrium binding constants were determined by plotting the average response values versus Fab concentration and fitting to a global one site-specific binding model in Prism (Graphpad).

Protein crystallization, data collection and processing

E104.v1.4DS S112F was crystallized at a concentration of 8 mg/mL via vapor diffusion in sitting well drops at 19°C in 0.2 M Na citrate and 20% PEG 3350. Crystals were cryoprotected in mother liquor supplemented with 10% glycerol. Diffraction data were collected at the Advanced Light Source (ALS) beamline 5.0.2. Data were processed to a resolution of 2.0 Å in XDS, and phases were obtained through molecular replacement with Phaser, using E104v1.WT variable and constant domains from a previously published crystal structure of the E104v1.WT-Tryptase complex (PDB: 6VVU, chains G and I) as search models.

E104.v1.4DS A114F was crystallized at a concentration of 10 mg/mL via vapor diffusion in sitting well drops at 19°C in 0.1 M Na citrate pH 4.5 and 20% PEG 4000. Crystals were cryoprotected in mother liquor supplemented with 10% glycerol. Diffraction data were collected at the Advanced Light Source (ALS) beamline 5.0.2. Data were processed to a resolution of 2.01 Å in XDS[5], and phases were obtained through molecular replacement with Phaser, using the crystal structure of E104v1.4DS S112F as the search model.

E104.v1.5DS and E104v1.6DS were crystallized at concentrations of 10 mg/mL and 7 mg/mL, respectively via vapor diffusion in hanging well drops at 19°C in 0.1 M Na Citrate pH 4.5 and 26% PEG 4000. Crystals were cryoprotected in mother liquor supplemented with 20% glycerol. Diffraction data were collected at Stanford Synchrotron Radiation Lightsource (SSRL) beamline 1.2.1. Data were processed to a resolution of 2.14 Å for E104.v1.5DS and 2.71 Å for E104.v1.6DS  in XDS, and phases were obtained through molecular replacement with Phaser, using the crystal structure of E104v1.4DS A114F as the search model.

Iterative rounds of model building and refinement were performed in COOT[6] and Phenix[7]. Figures were generated using PyMOL.

CryoEM sample preparation, data acquisition, and data processing

*Tryptase*

Tryptase-Fab complexes were prepared by incubating tetrameric tryptase with a 2-fold molar excess of Fab on ice for 30 min. The tryptase-Fab complex was then separated via size exclusion chromatography on a Superdex 200 3.2/300 or Superose 6 3.2/300 column (Cytiva) equilibrated in 20 mM MOPS pH 5.5, 800 mM NaCl. The peak fraction was subjected to mild crosslinking with 2.5 mM BS3 (ThermoFisher Scientific) at room temperature for 10 min. The crosslinking reaction was quenched by addition of 100mM Tris pH 7.5. Four µL of this sample were then applied to a grid, blotted in a Vitrobot MarkIV (ThermoFisher Scientific) at 4°C and 100% humidity, using a blotting time of 3 s and a blot force of 7, and plunge-frozen in liquid ethane cooled by liquid nitrogen. The tryptase-E104.v1.WT Fab complex was applied to a glow-discharged Quantifoil R0.6/1 Cu400 holey carbon grid (Electron Microscopy Sciences). The tryptase-E104.v1.2DS complex was applied to a glow-discharged Quantifoil R0.6/1 Au300 holey carbon grid (Electron Microscopy Sciences). The tryptase-E104.v1.4DS and tryptase-E104.v1.6DS complexes were applied to Quantifoil R0.6/1 Au300 holey carbon grids treated overnight with a thiol-reactive, self-assembling reaction mixture of 4 mM monothiolalkane(C11)PEG6-OH (11-mercaptoundecyl) hexaethyleneglycol (SPT-0011P6, SensoPath Technologies Inc., Bozeman, MT)[8]. Before application of the protein, the grids were removed from the SAM solution and rinsed with ethanol.

Movie stacks for tryptase-E104.v1.WT Fab were collected using SerialEM [9] on a Titan Krios (Thermo Fisher Scientific) operated at 300 kV and equipped with a BioQuantum energy filter operated with a 20 eV energy slit with a K2 Summit direct electron detector camera (Gatan). Images were recorded at a nominal magnification of 165,000x, corresponding to 0.824 Å per pixel. Each image stack contains 50 frames recorded every 0.2 s giving an accumulated dose of 54 e/Å$^2$ and a total exposure time of 10 s. Images were recorded with a set defocus range of 0.5 to 1.5 µm.

Movie stacks for tryptase-E104.v1.2DS, tryptase-E104.v1.4DS, and tryptase-E104.v1.6DS were collected using SerialEM[9] on a Titan Krios (Thermo Fisher Scientific) operated at 300 kV and equipped with a BioQuantum energy filter operated with a 20eV energy slit with a K3 Summit direct electron detector camera (Gatan). Images were recorded in EFTEM mode at a magnification of 105,000x corresponding to 0.838 Å per pixel, using a 20 eV energy slit. Each image stack contains 60 frames recorded every 0.05 s for an accumulated dose of ~65 e/Å$^2$ and a total exposure time of 3 s. Images were recorded with a set defocus range of 0.5 to 1.5 µm.

For all four tryptase-Fab complexes, image processing was performed as described in Fig. S4. Motion correction, CTF estimation, and particle picking were performed in cisTEM[10]. Particles were extracted from micrographs with CTF fit resolution less than 6 Å with a box size of 400 px and imported into Cryosparc. The particles were binned to a box size of 128 px and subjected to 2D classification to remove junk particles. The remaining particles were subjected to iterative rounds of multi-class ab initio reconstruction and heterogeneous refinement. The quality of the particles in the best classes was evaluated by running non-uniform refinement. The unbinned particles in the best class were then exported back into cisTEM. Prior to export, the Particle Sets tool in Cryosparc was used to randomly select a subset of the best particles from the 2DS, 4DS, and 6DS datasets in order to match the final number of particles in the WT dataset. In cisTEM, 3D refinement was performed using Auto-refine, using a low-pass filtered map from non-uniform refinement in Cryosparc as a reference. The auto-refined maps were then subjected to CTF refinement and manual refinement. The highest resolution used during refinement is indicated in Fig S4, panels D, I, N, and S. Local resolution maps were calculated using Relion[11].

The crystal structure of the tryptase-E104.v1 WT complex (PDB: 6VVU)[12] was used as an initial model to dock into the cryoEM maps in ChimeraX[13]. The resulting models were rebuilt and refined using COOT, ISOLDE[14], and Phenix. Sharpened maps were generated using the CryoEM module in COOT. Figures were generated using ChimeraX and PyMOL.

*Nav1.7*

Nav1.7-7A9 complexes were prepared by incubating Nav1.7 with a 1.2x molar excess of Fab at 4℃ for 30 minutes and separated on a Superose 6 3.2/300 column (Cytiva). Three µL from the peak fraction were applied to Quantifoil R2/2 Au300 holey carbon grids treated with a thiol-reactive, self-assembling reaction mixture of 4 mM monothiolalkane(C11)PEG6-OH (11-mercaptoundecyl) hexaethyleneglycol (SPT-0011P6, SensoPath Technologies Inc., Bozeman, MT)[8]. Before application of the protein, the grids were removed from the SAM solution and rinsed with ethanol. The grids were blotted in a Vitrobot MarkIV (ThermoFisher Scientific) at 4℃ and 100% humidity, using a blotting time of 3 s and a blot force of 7, and plunge-frozen in liquid ethane cooled by liquid nitrogen.

Movie stacks were collected using EPU on a Titan Krios (Thermo Fisher Scientific) operated at 300 kV and equipped with a Selectris energy filter and a Falcon4 detector. Images were recorded at a magnification of 165,000x corresponding to 0.731 Å per pixel, using a 20 eV energy slit. Each image stack contains 1077 frames recorded every 0.005 s for an accumulated dose of ~44 e/Å$^2$ and a total exposure time of 5 s. Images were recorded with a set defocus range of 0.5 to 1.5 µm.

All image processing was performed in CryoSPARC[15], as summarized in Fig. S5. Patch motion correction, patch CTF estimation, and particle picking were performed using CryoSPARC Live. Micrographs with CTF fit resolutions worse than 6.5 Å were rejected. The blob picker was used with a minimum radius of 160 Å and maximum radius of 220 Å. Particles were extracted with a box size of 512 px and binned to 128 px. 2D classification on a small subset of the data was used to identify 2D classes for generation of initial model via ab initio reconstruction. This model was used to generate templates for particle picking with a radius of 220 Å. The resulting particle stack was subjected to 2D classification to remove junk particles. The remaining particles were subjected to iterative rounds of multi-class ab initio reconstruction and heterogeneous refinement. The quality of the particles in the best classes were evaluated by running non-uniform refinement. The particles from the best classes were subjected to 3D classification into 10 classes with a mask around Nav1.7 and the Fab variable domain. Classes where the 4-helix bundle was best-resolved were selected and re-extracted with a box size of 512 px and binned to

400 px. Non-uniform refinement of the re-extracted particles led to a consensus 3D reconstruction. Local refinement with a mask around both Fabs was used to improve alignments of the Fabs prior to particle subtraction to remove the Fabs. The signal-subtracted particles were then subjected to local refinement with a mask around Nav1.7. Phenix.combine_focused_maps was used to generate a composite map using the consensus map and local refinement maps focused on the Fabs and Nav1.7 as inputs. The local resolution of the composite map was calculated using the Local Resolution job in Phenix with the composite half maps produced by Phenix.combine_focused_maps.

Previously published structures of the Nav1.7-ProTx2-7A9.WT complex (PDB: 6N4Q)[16] and Nav1.7-NavAb (PDB: 5EK0)[2] were used as an initial models for the 7A9 Fab and Nav1.7, respectively, to dock into the cryoEM maps in ChimeraX. The resulting models were rebuilt and refined using COOT, ISOLDE, and Phenix. Sharpened maps were generated using the CryoEM module in COOT. Figures were generated using ChimeraX and PyMOL.

*CD20*

CD20-RTX complexes were prepared by incubating CD20 with a 1.2 molar excess of Fab at 4℃ for 30 minutes and separated on a Superose 6 3.2/300 column (Cytiva). 3 µL from the peak fraction were applied to Quantifoil R0.6/1 Au300 holey carbon grids treated overnight with a thiol-reactive, self-assembling reaction mixture of 4 mM monothiolalkane(C11)PEG6-OH (11-mercaptoundecyl) hexaethyleneglycol (SPT-0011P6, SensoPath Technologies Inc., Bozeman, MT)[8]. Before application of the protein, the grids were removed from the SAM solution and rinsed with ethanol. The grids were blotted in a Vitrobot MarkIV (ThermoFisher Scientific) at 4℃ and 100% humidity, using a blotting time of 5 s and a blot force of 8, and plunge-frozen in liquid ethane cooled by liquid nitrogen.

Movie stacks for CD20-RTX.WT were collected from 1 grid using EPU on a Titan Krios (Thermo Fisher Scientific) operated at 300 kV and equipped with a Selectris energy filter and a Falcon4 detector. Images were recorded at a magnification of 165,000x corresponding to 0.731 Å per pixel, using a 20 eV energy slit. Each image stack contains 1011 frames recorded every 0.004 s for an accumulated dose of 37 e/Å$^2$ and a total exposure time of 4 s. Images were recorded with a set defocus range of 0.5 to 1.5 µm.

Movie stacks for CD20-RTX.4DS were collected from 4 grids using EPU on a Titan Krios (Thermo Fisher Scientific) operated at 300 kV and equipped with a Selectris energy filter and a Falcon4 detector. Images were recorded at a magnification of 165,000x corresponding to 0.731 Å per pixel, using a 20 eV energy slit. Each image stack contains 1001 frames recorded every 0.005 s for an accumulated dose of 45 e/Å$^2$ and a total exposure time of 5 s. Images were recorded with a set defocus range of 0.5 to 1.5 µm.

All image processing was performed in CryoSPARC, as summarized in Fig. S6. Patch motion correction, patch CTF estimation, and particle picking were performed using CryoSPARC Live. Micrographs with CTF fit resolutions worse than 4.0 Å were rejected. The blob picker was used with a minimum radius of 150 Å and maximum radius of 220 Å. Particles were extracted with a box size of 512 px and binned to 128 px. 2D classification on a small subset of the data was used to identify 2D classes for generation of initial model via ab initio reconstruction. This model was used to generate templates for particle picking with a radius of 200 Å. The resulting particle stack was subjected to 2D classification to remove junk particles. The remaining particles were subjected to iterative rounds of multi-class ab initio reconstruction and heterogeneous refinement. The quality of the particles in the best classes were evaluated by running non-uniform refinement. These particles from the best classes were re-extracted with a box size of 512 px and binned to 400 px. Non-uniform refinement of the re-extracted particles led to a consensus 3D reconstruction. Local refinement was performed with a mask around the variable domains of both Fabs and CD20. The fulcrum for the local refinement was also shifted to the center of mass for the CD20 portion of the structure. The resulting volume was used as input for another local refinement with a mask around only CD20. Phenix.combine_focused_maps was used to generate a composite map using the consensus map and local refinement maps as inputs. The local resolution of the composite map was calculated using the Local Resolution job in Phenix with the composite half maps produced by Phenix.combine_focused_maps.

The model for the structure of the CD20-RTX.WT complex (PDB: 6VJA)[3] was used as an initial model to dock into the cryoEM maps in ChimeraX. The resulting models were rebuilt and refined using COOT, ISOLDE, and Phenix. Sharpened maps were generated using the CryoEM module in COOT. Figures were generated using ChimeraX and PyMOL.

*Ang2*

Ang2 was incubated with an equimolar amount of the 5A12.6DS Fab and a 2-fold molar excess of ProA-ProG fusion protein on ice for 30 min. This mixture was injected onto a Superdex 200 3.2/300 column equilibrated in 20 mM HEPES pH 7.5, 150 mM NaCl. The peak fraction was subjected to mild crosslinking with 0.5 mM BS3 at room temperature for 10 min. The crosslinking reaction was quenched by addition of 100mM Tris pH 7.5. 4 µL of this sample were then applied to glow-discharged Quantifoil R0.6/1 Au300 holey carbon grids, blotted in a Vitrobot MarkIV at 4°C and 100% humidity using a blotting time of 6 s and a blot force of 7, and plunge-frozen in liquid ethane cooled by liquid nitrogen.

Movie stacks were collected using SerialEM[9] on a Titan Krios (Thermo Fisher Scientific) operated at 300 kV and equipped with a BioQuantum energy filter operated with a 20eV energy slit with a K3 Summit direct electron detector camera (Gatan). Images were recorded in EFTEM mode at a magnification of 105,000x corresponding to 0.838 Å per pixel, using a 20 eV energy slit. Each image stack contains 119 frames recorded every 0.05 s for an accumulated dose of 69 e/$Å^2$ and a total exposure time of 6 s. Images were recorded with a set defocus range of 0.5 to 1.5 µm.

All image processing was performed using CryoSPARC as summarized in the schematic in Fig. S7. Patch motion correction, patch CTF estimation, and particle picking were performed using CryoSPARC Live. Micrographs with CTF fit resolutions worse than 3.7 Å or with relative ice thickness higher than 1.077 were rejected. The blob picker was used with a minimum radius of 50 Å and maximum radius of 150 Å, using both a circular blob and elliptical blob, as well as a minimum separation distance of 0.3 diameters. Particles were extracted with a box size of 400 px and binned to 128 px. 2D classification of a small subset of the data was performed to identify

2D classes for generation of initial model via ab initio reconstruction. This model was used to generate templates for particle picking with a radius of 100 Å. The resulting particle stack was subjected to 2D classification to remove junk particles. The remaining particles were subjected to iterative rounds of multi-class ab initio reconstruction and heterogeneous refinement, resulting in a stack of 1,017,611 particles, which were re-extracted with a box size of 400 px and binned to 324 px. These particles were then subjected to non-uniform refinement, resulting in a 3D reconstruction with a GS-FSC resolution of 2.69 Å. A local resolution map was calculated using the Local Resolution Estimation job in CryoSPARC.

The crystal structure of the Ang2-5A12.WT complex (PDB: 4ZFG)[4] was used as an initial model to dock into the cryoEM map in ChimeraX. The resulting model was rebuilt and refined using COOT, ISOLDE, and Phenix. Sharpened maps were generated using the CryoEM module in COOT. Figures were generated using ChimeraX and PyMOL.

*$KRAS^{G12C}$*

$KRAS^{G12C}$ was covalently modified with GNE-1952 as previously described[17]. $KRAS^{G12C}$ was incubated for 4 hrs at room temperature with 150 μM GNE-1952, 5 mM GDP, and 20 mM EDTA. Complete covalent modification was confirmed via mass spectrometry. The modified protein was then buffer exchanged into 20 mM HEPES pH 7.0, 100 mM NaCl, 1 mM $MgCl_2$, 5 μM GDP through size exclusion chromatography on a Superdex 75 16/60 column (Cytiva). The $KRAS^{G12C}$-GNE-1952 adduct was incubated with a 2-fold molar excess of the 2H11.4DS Fab on ice for 30 min. This mixture was then injected onto a Superdex 200 3.2/300 column equilibrated in 20 mM HEPES pH 7.0, 100 mM NaCl, 1 mM $MgCl_2$, 5 μM GDP. The peak fraction was subjected to mild crosslinking with 0.5 mM BS3 at room temperature for 10 min. 4 μL of this sample were then applied to glow-discharged Quantifoil R0.6/1 Au300 holey carbon grids, blotted in a Vitrobot MarkIV at 4°C and 100% humidity using a blotting time of 6 s and a blot force of 7, and plunge-frozen in liquid ethane cooled by liquid nitrogen.

Movie stacks were collected using EPU on a Titan Krios (Thermo Fisher Scientific) operated at 300 kV and equipped with a Selectris energy filter and a Falcon4 detector. Images were recorded at a magnification of 165,000x corresponding to 0.731 Å per pixel, using a 20 eV energy slit. Each image stack contains 1077 frames recorded every 0.005 s for an accumulated dose of 40 $e/Å^2$ and a total exposure time of 2.2 s. Images were recorded with a set defocus range of 0.5 to 1.5 μm.

All image processing was performed using CryoSPARC as summarized in the schematic in Fig. S8. Patch motion correction, patch CTF estimation, and particle picking were performed using CryoSPARC Live. Micrographs with CTF fit resolutions worse than 4.0 Å or with relative ice thickness higher than 1.12 were rejected. The blob picker was used with a minimum radius of 50 Å and maximum radius of 150 Å, using both a circular blob and elliptical blob, as well as a minimum separation distance of 0.5 diameters. Particles were extracted with a box size of 324 px and binned to 128 px. 2D classification of a small subset of the data was performed to identify 2D classes for generation of initial model via ab initio reconstruction. This model was used to generate templates for particle picking with a radius of 150 Å. The resulting particle stack was subjected to 2D classification to remove junk particles. The remaining particles were subjected to iterative rounds of multi-class ab initio reconstruction and heterogeneous refinement, resulting in a stack of 926,738 particles, which were re-extracted with a box size of 324 px and binned to 224 px. These particles were then subjected to non-uniform refinement, including refinement of per-particle defocus and per-group CTF parameters (tilt and trefoil), resulting in a 3D reconstruction with a GS-FSC resolution of 2.83 Å. Local refinement was then performed using a mask around KRAS and the 2H11.4DS variable domain, leading to a 3D reconstruction with a GS-FSC resolution of 2.74 Å. Local resolution maps were calculated using the Local Resolution Estimation job in CryoSPARC.

The crystal structure of the KRAS$^{G12C}$-GNE-1952-2H11.WT complex (PDB: 7RP3)[17] was used as an initial model to dock into the cryoEM maps in ChimeraX. The resulting model was rebuilt and refined using COOT, ISOLDE, and Phenix. Sharpened maps were generated using the CryoEM module in COOT. Figures were generated using ChimeraX and PyMOL.

### Extended Data

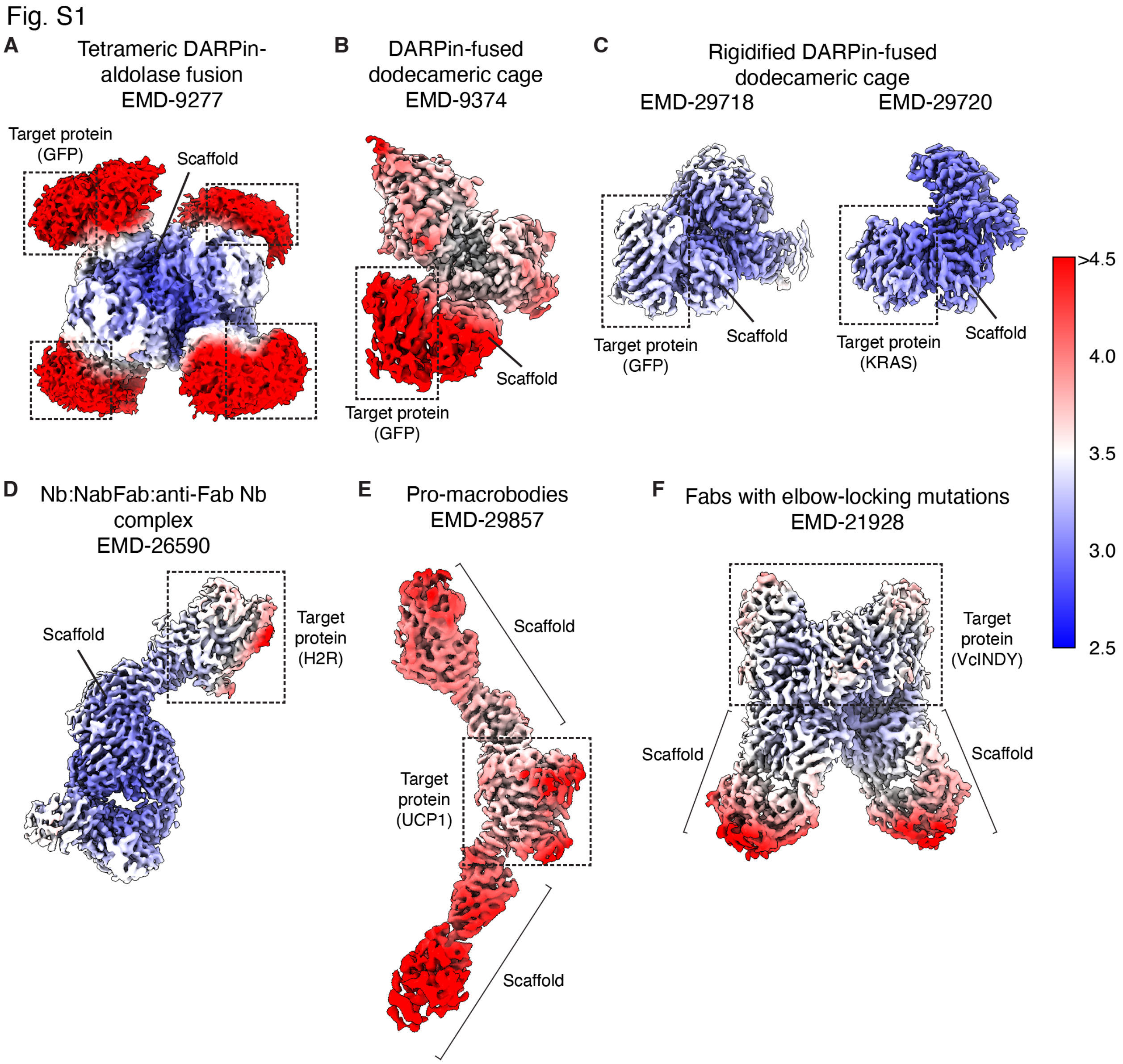

**Extended Data Fig. 1: Flexibility in previously published structure chaperones.**
**A.** Local resolution map of a tetrameric darpin-aldolase fusion bound to target protein GFP[18]. The aldolase scaffold is much more rigid and resolved to a higher resolution than the DARPin or GFP, likely owing to flexibility of the linker between the DARPin and aldolase, as well as flexibility within the DARPin itself. **B.** Local resolution map of one subunit of a dodecameric cage fused to a DARPin binding the target protein GFP[19]. The resolution of GFP is much lower than for the cage proteins, likely due to flexibility in the linker between the cage subunit and DARPin, as well as flexibility of the DARPin itself. **C.** Local resolution maps of GFP (left) and KRAS (right) bound to a subunit of a DARPin-dodecameric cage fusion in which the DARPins

are fused more rigidly to the cage than in (C)[20]. The increase in rigidity is apparent in comparison with (C), but there is still some loss of resolution in GFP and KRAS distal from the DARPin, indicating some remaining flexibility in the scaffold. **D.** Local resolution map of histamine receptor 2 (H2R) bound to a scaffold made up of an anti-H2R nanobody (Nb):anti-Nb Fab (NabFab):anti-Fab Nb complex[21,22]. While the Nb:NabFab core of the scaffold appears rigid and well-resolved, the target H2R has lower resolution. **E.** Local resolution map of mitochondrial uncoupling protein 1 (UCP1) bound to two pro-macrobodies, PMb65 and PMb71, consisting of Nbs fused to maltose binding protein (MBP) with a di-proline linker[23,24]. For both pro-macrobodies, the Nb portion of the scaffold is better resolved than the MBP portion, as well as the target protein, indicating flexibility in the scaffold. **F.** Local resolution map of a homodimer of *V. cholerae* $Na^+$-dependent dicarboxylate transporter VcINDY bound to two copies of a Fab in which the heavy chain has been mutated to reduce elbow flexibility[25,26]. Despite the "elbow-locking" mutations, the low resolution of the constant domains relative to the variable domains of the Fabs indicates that the Fab is still very flexible. Local resolution maps for (A-F) were calculated using Relion and the half-maps deposited for each structure in EMDB.

Fig. S2

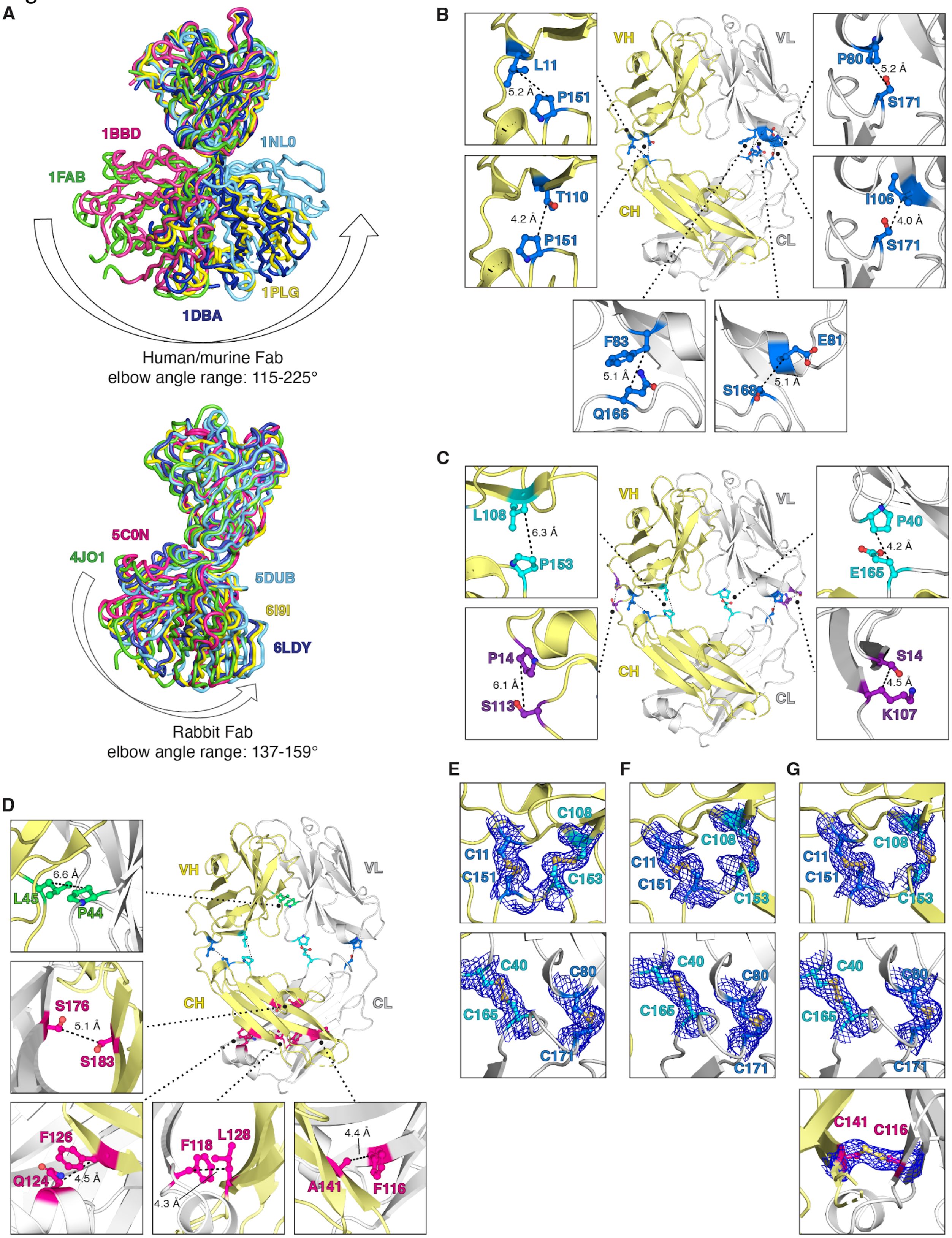

A
1BBD
1NL0
1FAB
1PLG
1DBA
Human/murine Fab
elbow angle range: 115-225°
5C0N
4JO1
5DUB
6I9I
6LDY
Rabbit Fab
elbow angle range: 137-159°
B
VH
VL
CH
CL
L11
5.2 Å
P151
T110
4.2 Å
P151
P80
5.2 Å
S171
I106
4.0 Å
S171
F83
5.1 Å
Q166
E81
S168
5.1 Å
C
L108
6.3 Å
P153
P14
6.1 Å
S113
P40
4.2 Å
E165
S14
4.5 Å
K107
D
L45
6.6 Å
P44
S176
5.1 Å
S183
F126
Q124
4.5 Å
F118
L128
4.3 Å
4.4 Å
A141
F116
E
F
G
C11
C151
C108
C153
C40
C165
C80
C171
C141
C116

**Extended Data Fig. 2: Design of conformationally rigid Fabs.**

**A.** Top, VH-VL-based alignment illustrating the wide range of elbow angles adopted by selected human, murine, and rat Fabs (adapted from Stanfield et al[27]). Fab PDB IDs: 1BBD (magenta), 1FAB (green), 1DBA (dark blue), 1PLG (yellow), 1NL0 (light blue). Bottom, VH-VL-based alignment illustrating the narrow range of elbow angles adopted by rabbit Fabs. PDB IDs: 5C0N (magenta), 4JO1 (green), 6LDY (dark blue), 6I9I (yellow), 5DUB (light blue). **B-D.** Residues mutated to introduce intrachain disulfides in the elbow region (B and C), as well as interchain disulfides in the variable and constant domains (D) mapped onto the crystal structure of the E104.v1.WT Fab (PDB: 6VVU) shown in cartoon representation with the heavy chain in yellow and light chain in light gray. Insets show zoomed-in views of pairs of mutations and the Cβ-Cβ distances (dashed lines) between these residues. Distances were measured in PyMOL. Fabs containing the following pairs of residues mutated to cysteines resulted in loss of expression: $P14^{HC}$:$S113^{HC}$, $S14^{LC}$:$K107^{LC}$, $L45^{HC}$:$P44^{LC}$, $S176^{HC}$:$S183^{LC}$. **E-G.** Electron density contoured at 1σ from crystal structures of E104.v1.4DS.S112F (E), E104.v1.4DS.A114F (F), and E104.v1.5DS.A114F (G), confirming proper formation of the engineered disulfides in these constructs. Mutations S112F and A114F were not necessary for rigidity of the Fab, but enabled crystallization of the various Rigid Fab designs.

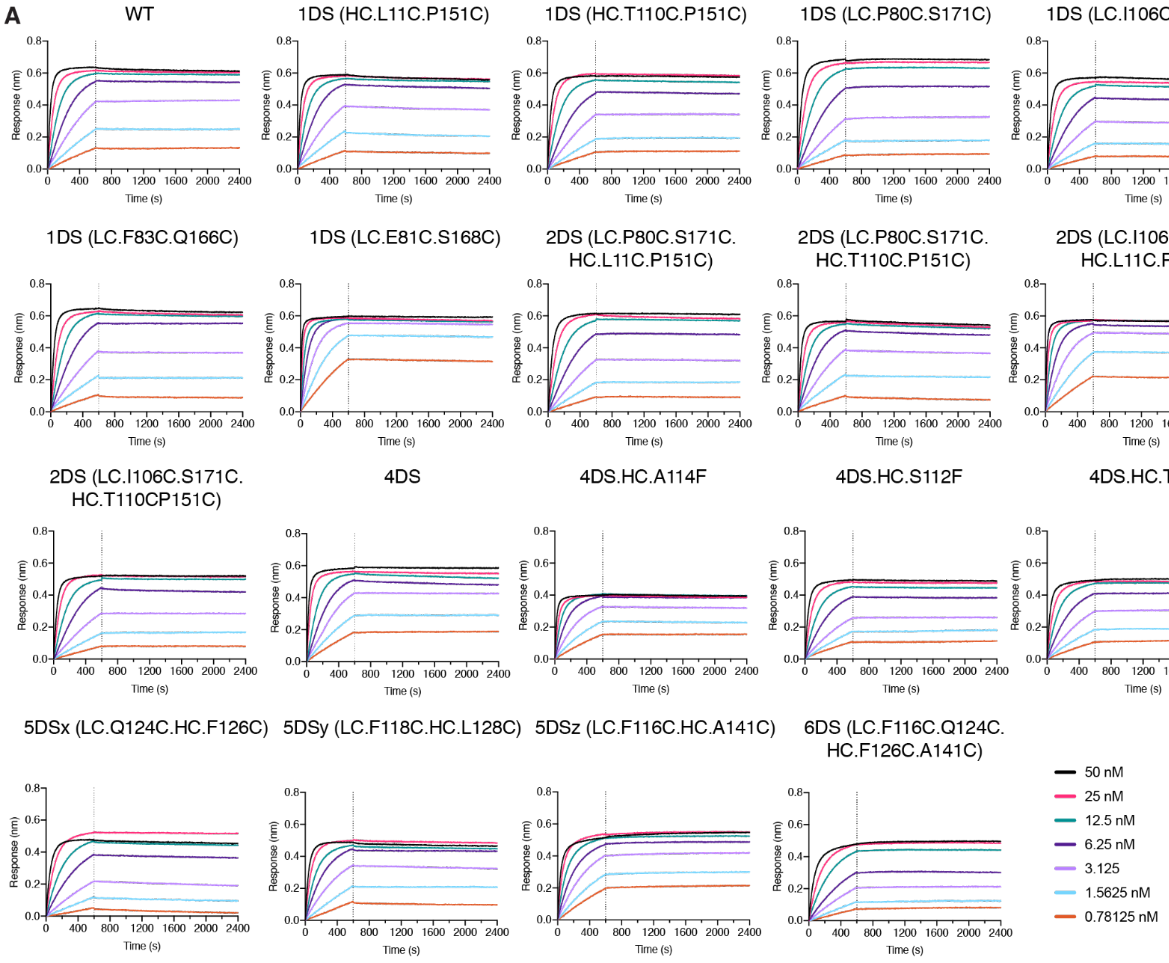

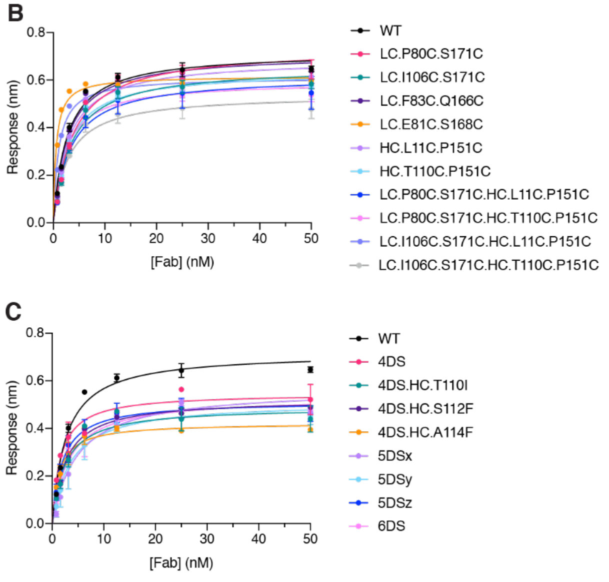

**D**

| Construct | Steady State Kd (nM) |
|---|---|
| WT | 2.65 |
| 1DS (LC.P80C.S171C) | 3.75 |
| 1DS (LC.I106C.S171C) | 3.41 |
| 1DS (LC.F83C.Q166C) | 2.74 |
| 1DS (LC.E81C.S168C) | 0.55 |
| 1DS (HC.L11C.P151C) | 2.87 |
| 1DS (HC.T110C.P151C) | 2.82 |
| 2DS (LC.P80C.S171C.HC.L11C.P151C) | 3.25 |
| 2DS (LC.P80C.S171C.HC.T110C.P151C) | 2.4 |
| 2DS (LC.I106C.S171C.HC.L11C.P151C) | 1.05 |
| 2DS (LC.I106C.S171C.HC.T110C.P151C) | 2.87 |
| 4DS | 1.63 |
| 4DS.HC.A114F | 1.37 |
| 4DS.HC.T110I | 2.46 |
| 4DS.HC.S112F | 2.59 |
| 5DSx (LC.Q124C.HC.F126C) | 4.81 |
| 5DSy (LC.F118C.HC.L128C) | 3.51 |
| 5DSz (LC.F116C.HC.A141C) | 1.94 |
| 6DS (LC.F116C.Q124C.HC.F126C.A141C) | 3.16 |

**Extended Data Fig.3: Rigid Fab mutations do not affect antigen binding.**
**A.** Representative biolayer interferometry (BLI) traces showing binding of a dilution series of the indicated E104.v1 Fab constructs to immobilized biotinylated tryptase. Sensorgrams were

normalized to a reference well containing only buffer. The dashed line indicates the beginning of the dissociation step. **B and C.** BLI equilibrium values from panel A are plotted as a function of Fab concentration. Circles and error bars represent average response ± SEM. Fitting of these data with a 1:1 binding model was used to calculate the apparent $K_d$ values shown in (D).
**D.** Steady state $K_d$ values measured in tryptase-Fab BLI experiments.

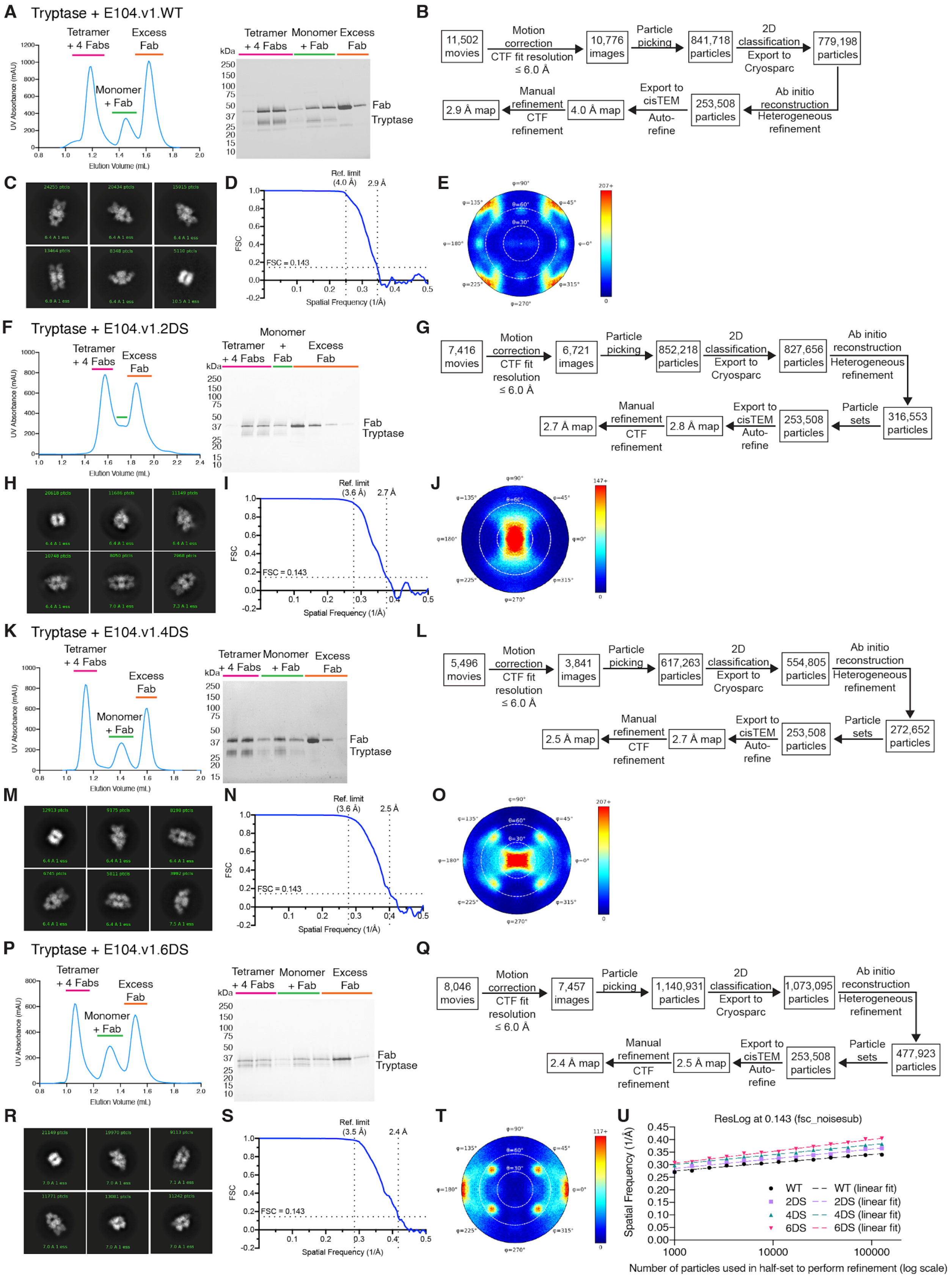
A
Tryptase + E104.v1.WT
Tetramer + 4 Fabs
Excess Fab
Monomer + Fab
UV Absorbance (mAU)
Elution Volume (mL)
kDa
Tetramer + 4 Fabs
Monomer + Fab
Excess Fab
Fab
Tryptase
B
11,502 movies
Motion correction
CTF fit resolution ≤ 6.0 Å
10,776 images
Particle picking
841,718 particles
2D classification
Export to Cryosparc
779,198 particles
Ab initio reconstruction
Heterogeneous refinement
253,508 particles
Export to cisTEM
Auto-refine
4.0 Å map
Manual refinement
CTF refinement
2.9 Å map
C
D
Ref. limit (4.0 Å)
2.9 Å
FSC
FSC = 0.143
Spatial Frequency (1/Å)
E
F
Tryptase + E104.v1.2DS
Tetramer + 4 Fabs
Excess Fab
UV Absorbance (mAU)
Elution Volume (mL)
Monomer + Fab
kDa
Fab
Tryptase
G
7,416 movies
Motion correction
CTF fit resolution ≤ 6.0 Å
6,721 images
Particle picking
852,218 particles
2D classification
Export to Cryosparc
827,656 particles
Ab initio reconstruction
Heterogeneous refinement
316,553 particles
Particle sets
253,508 particles
Export to cisTEM
Auto-refine
2.8 Å map
Manual refinement
CTF refinement
2.7 Å map
H
I
Ref. limit (3.6 Å)
2.7 Å
FSC
FSC = 0.143
Spatial Frequency (1/Å)
J
K
Tryptase + E104.v1.4DS
Tetramer + 4 Fabs
Excess Fab
Monomer + Fab
UV Absorbance (mAU)
Elution Volume (mL)
kDa
Fab
Tryptase
L
5,496 movies
Motion correction
CTF fit resolution ≤ 6.0 Å
3,841 images
Particle picking
617,263 particles
2D classification
Export to Cryosparc
554,805 particles
Ab initio reconstruction
Heterogeneous refinement
272,652 particles
Particle sets
253,508 particles
Export to cisTEM
Auto-refine
2.7 Å map
Manual refinement
CTF refinement
2.5 Å map
M
N
Ref. limit (3.6 Å)
2.5 Å
FSC
FSC = 0.143
Spatial Frequency (1/Å)
O
P
Tryptase + E104.v1.6DS
Tetramer + 4 Fabs
Excess Fab
Monomer + Fab
UV Absorbance (mAU)
Elution Volume (mL)
kDa
Fab
Tryptase
Q
8,046 movies
Motion correction
CTF fit resolution ≤ 6.0 Å
7,457 images
Particle picking
1,140,931 particles
2D classification
Export to Cryosparc
1,073,095 particles
Ab initio reconstruction
Heterogeneous refinement
477,923 particles
Particle sets
253,508 particles
Export to cisTEM
Auto-refine
2.5 Å map
Manual refinement
CTF refinement
2.4 Å map
R
S
Ref. limit (3.5 Å)
2.4 Å
FSC
FSC = 0.143
Spatial Frequency (1/Å)
T
U
ResLog at 0.143 (fsc_noisesub)
Spatial Frequency (1/Å)
WT
WT (linear fit)
2DS
2DS (linear fit)
4DS
4DS (linear fit)
6DS
6DS (linear fit)
Number of particles used in half-set to perform refinement (log scale)

**Extended Data Fig.4: CryoEM sample preparation and image processing for tryptase-E104.v1 Fab complexes.**

**A.** SEC elution profile and gel showing formation of tryptase-E104.v1.WT Fab complex. **B-E.** Representative 2D classes (B), image processing workflow (C), FSC curve (D), and angular distribution plot (E) for tryptase-E104.v1.WT dataset. **F.** SEC elution profile and gel showing formation of tryptase-E104.v1.2DS Fab complex. **G-J.** Representative 2D classes (G), image processing workflow (H), FSC curve (I), and angular distribution plot (J) for tryptase-E104.v1.2DS dataset. **K.** SEC elution profile and gel showing formation of tryptase-E104.v1.4DS Fab complex. **L-O.** Representative 2D classes (L), image processing workflow (M), FSC curve (N), and angular distribution plot (O) for tryptase-E104.v1.4DS dataset. **P.** SEC elution profile and gel showing formation of tryptase-E104.v1.6DS Fab complex. **Q-T.** Representative 2D classes (Q), image processing workflow (R), FSC curve (S), and angular distribution plot (T) for tryptase-E104.v1.6DS dataset. **U.** ResLog plot ($FSC_{0.143}$) comparing the WT, 2DS, 4DS, and 6DS datasets. fsc_noisesub was calculated for each reconstruction using the ResLog Analysis job in CryoSparc.

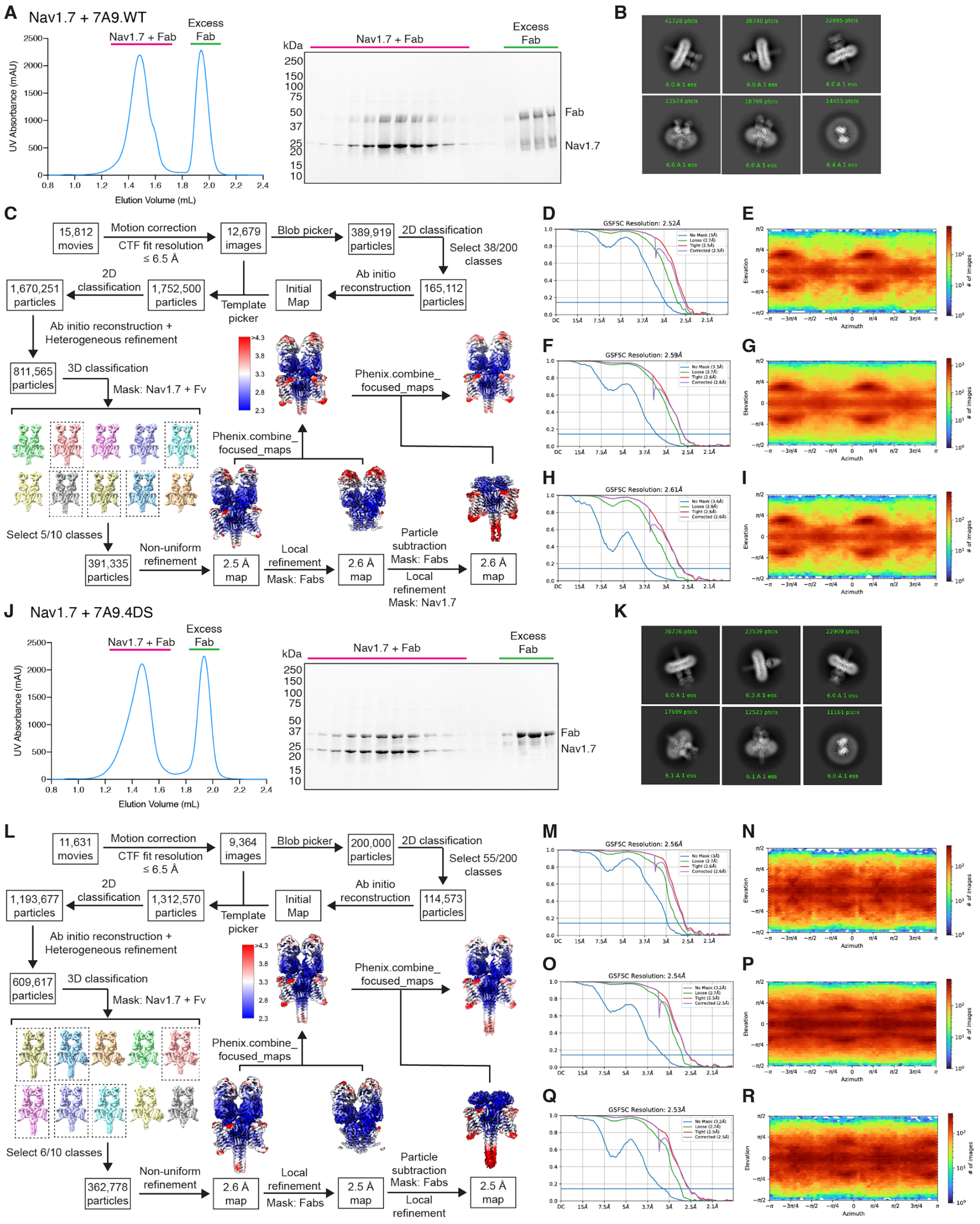

**Extended Data Fig. 5: CryoEM sample preparation and image processing for Nav-7A9 Fab complexes.**

**A.** SEC elution profile and gel showing formation of Nav1.7-7A9.WT Fab complex. **B-E.** Representative 2D classes (B), image processing workflow (C), FSC curve (D) and angular distribution plot (E) for consensus refinement, FSC curve (F) and angular distribution plot (G) for local refinement with mask around Fabs, FSC curve (H) and angular distribution plot (I) for local refinement with mask around Nav1.7 following particle subtraction for Nav1.7-7A9.WT dataset. **J.** SEC elution profile and gel showing formation of Nav1.7-7A9.4DS Fab complex. **K-R.** Representative 2D classes (K), image processing workflow (L), FSC curve (M) and angular distribution plot (N) for consensus refinement, FSC curve (O) and angular distribution plot (P) for local refinement with mask around Fabs, FSC curve (Q) and angular distribution plot (R) for local refinement with mask around Nav1.7 following particle subtraction for Nav1.7-7A9.4DS dataset.

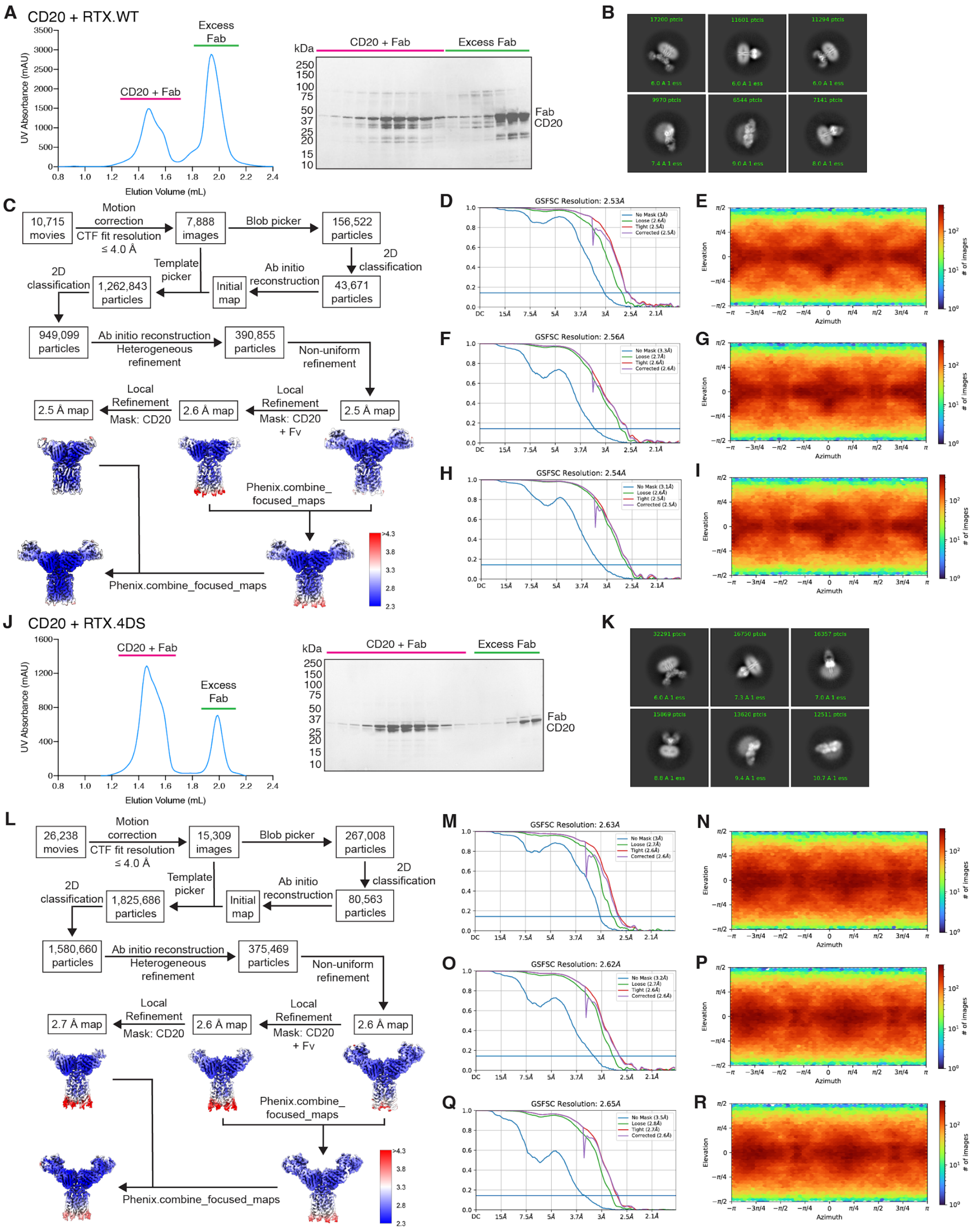

**Extended Data Fig. 6: CryoEM sample preparation and image processing for CD20-RTX Fab complexes.**

**A.** SEC elution profile and gel showing formation of CD20.RTX Fab complex. **B-I.** Representative 2D classes (B), image processing workflow (C), FSC curve (D) and angular distribution plot (E) for consensus refinement, FSC curve (F) and angular distribution plot (G) for local refinement with mask around CD20 and variable domains, FSC curve (H) and angular distribution plot (I) for local refinement with mask around CD20 for CD20-RTX.WT dataset. **J.** SEC elution profile and gel showing formation of CD20.RTX Fab complex. **K-R.** Representative 2D classes (K), image processing workflow (L), FSC curve (M) and angular distribution plot (N) for consensus refinement, FSC curve (O) and angular distribution plot (P) for local refinement with mask around CD20 and variable domains, FSC curve (Q) and angular distribution plot (R) for local refinement with mask around CD20 for CD20-RTX.WT dataset.

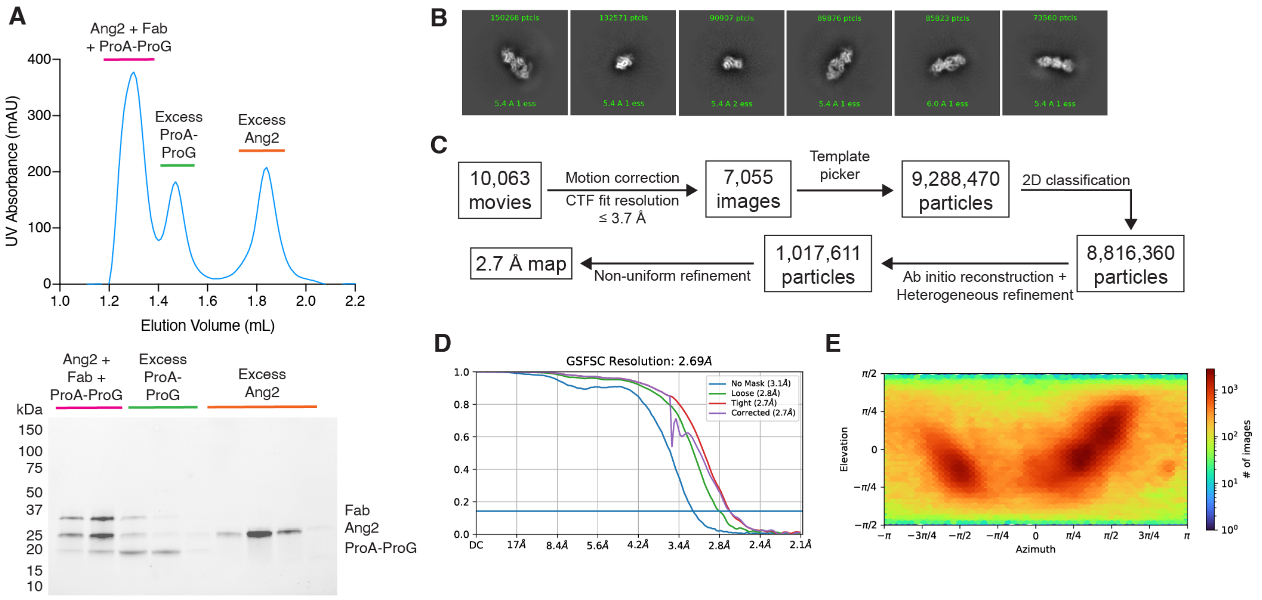

**Extended Data Fig.7: CryoEM sample preparation and image processing for Ang2-5A12.6DS Fab complex.**

**A.** SEC elution profile and gel showing formation of Ang2-ProA-ProG-5A12.6DS Fab complex. **B-E.** Representative 2D classes (B), image processing workflow (C), FSC curve (D), and angular distribution plot (E) for Ang2-5A12.6DS dataset.

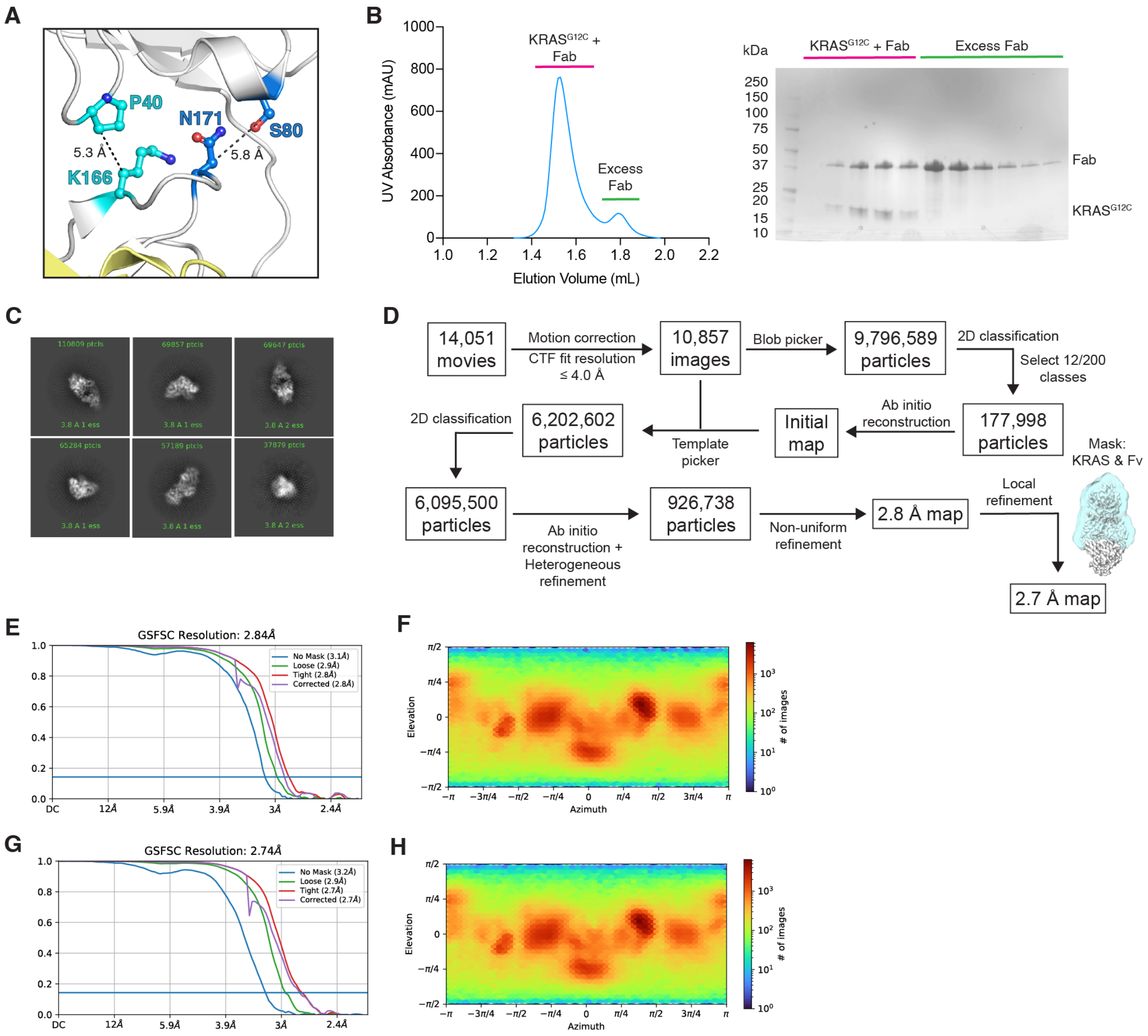

**Extended Data Fig. 8: CryoEM sample preparation and image processing for KRAS$^{G12C}$-GNE-1952-2H11.4DS Fab complex.**

**A.** Design of LC elbow disulfide mutations for 2H11 Fab. Mutated residues are mapped onto a cartoon representation of 2H11 LC (light gray) from a crystal structure of a KRAS$^{G12C}$-2H11 crystal structure (PDB:7RP2), with Cβ-Cβ distances shown in dashed lines. Since P80 and N171 (blue) are too far to enable disulfide mutation by mutating both these sites to cysteine, a cysteine was inserted between N170 and N171. The other LC elbow disulfide was formed using cysteine mutations at P40 and K166 (cyan). **B.** SEC elution profile and gel showing formation of KRAS$^{G12C}$-GNE-1952-2H11.4DS Fab complex. **C-H.** Representative 2D classes (C), image processing workflow (D), FSC curve (E) and angular distribution plot (F) for consensus refinement, FSC curve (G) and angular distribution plot (H) for local refinement with mask around KRAS and the Fab variable domain for the KRAS$^{G12C}$-GNE-1952-2H11.4DS dataset.

**A**

```
E104v1_WT      DIQMTQSPSSLSASVGDRVTITCQSIK-SVYNNRLGWYQQKPGKAPKLLIYETSILTSGVPSRFSGSGSGTDFTLTISSLQPEDFATYYCAGGFDRSGDTTFGQGTKVEIK-RTVAAPSV    118
E104v1_2DS     DIQMTQSPSSLSASVGDRVTITCQSIK-SVYNNRLGWYQQKPGKAPKLLIYETSILTSGVPSRFSGSGSGTDFTLTISSLQCEDFATYYCAGGFDRSGDTTFGQGTKVEIK-RTVAAPSV    118
E104v1_4DS     DIQMTQSPSSLSASVGDRVTITCQSIK-SVYNNRLGWYQQKCGKAPKLLIYETSILTSGVPSRFSGSGSGTDFTLTISSLQCEDFATYYCAGGFDRSGDTTFGQGTKVEIK-RTVAAPSV    118
E104v1_6DS     DIQMTQSPSSLSASVGDRVTITCQSIK-SVYNNRLGWYQQKCGKAPKLLIYETSILTSGVPSRFSGSGSGTDFTLTISSLQCEDFATYYCAGGFDRSGDTTFGQGTKVEIK-RTVAAPSV    118
7A9_WT         EIVLTQSPALMAASPGEKVTITCSVSL-SISSSNLFWYQQKSETSPKPWIYGTSKLASGVPVRFSGSGSGTSYSLTISSMEAEDAATYYCQQWSS--HSFTFGGGTKLEIK-RADAAPTV    116
7A9_4DS        EIVLTQSPALMAASPGEKVTITCSVSL-SISSSNLFWYQQKCETSPKPWIYGTSKLASGVPVRFSGSGSGTSYSLTISSMECEDAATYYCQQWSS--HSFTFGGGTKLEIK-RADAAPTV    116
RTX_WT         QIVLSQSPAILSASPGEKVTMTCRASS-SVS--YIHWFQQKPGSSPKPWIYATSNLASGVPVRFSGSGSGTSYSLTISRVEAEDAATYYCQQWT--SNPPTFGGGTKLEIK-RTVAAPSV    114
RTX_4DS        QIVLSQSPAILSASPGEKVTMTCRASS-SVS--YIHWFQQKCGSSPKPWIYATSNLASGVPVRFSGSGSGTSYSLTISRVECEDAATYYCQQWT--SNPPTFGGGTKLEIK-RTVAAPSV    114
5A12_WT        DIQMTQSPSSLSASVGDRVTITCRASQ-FLSSFGVAWYQQKPGKAPKLLIYGASSLYSGVPSRFSGSGSGTDFTLTISSLQPEDFATYYCQQGLL--SPLTFGQGTKVEIK-RTVAAPSV    116
5A12_6DS       DIQMTQSPSSLSASVGDRVTITCRASQ-FLSSFGVAWYQQKCGKAPKLLIYGASSLYSGVPSRFSGSGSGTDFTLTISSLQCEDFATYYCQQGLL--SPLTFGQGTKVEIK-RTVAAPSV    116
2H11_WT        --SVLTQPPSASGTPGQRVTISCSGSSSNIGSNYVYWYQQLPGTAPKLLIYRNNQRPSGVPDRFSGSKSGTSASLAISGLRSEDEADYYCAAWDERLSGWVFGGGTKLTVLGQPKAAPSV    118
2H11_4DS       --SVLTQPPSASGTPGQRVTISCSGSSSNIGSNYVYWYQQLCGTAPKLLIYRNNQRPSGVPDRFSGSKSGTSASLAISGLRCEDEADYYCAAWDERLSGWVFGGGTKLTVLGQPKAAPSV    118
                   :  .*   :.: *::**::*       :     : *:**    .:**  **  .    **** ***** ***. :*:** :. ** * ***           .** ***: :  :  ***:*

E104v1_WT      FIFPPSDEQLKSGTASVVCLLNNFYPREAKVQWKVDNALQSGNSQESVTEQDSKDSTYSLSSTLTLSKADYEKHKVYACEVTHQGLSSPVTKSFNRGEC-    217
E104v1_2DS     FIFPPSDEQLKSGTASVVCLLNNFYPREAKVQWKVDNALQSGNSQESVTEQDSKDCTYSLSSTLTLSKADYEKHKVYACEVTHQGLSSPVTKSFNRGEC-    217
E104v1_4DS     FIFPPSDEQLKSGTASVVCLLNNFYPREAKVQWKVDNALQSGNSQESVTCQDSKDCTYSLSSTLTLSKADYEKHKVYACEVTHQGLSSPVTKSFNRGEC-    217
E104v1_6DS     CIFPPSDECLKSGTASVVCLLNNFYPREAKVQWKVDNALQSGNSQESVTCQDSKDCTYSLSSTLTLSKADYEKHKVYACEVTHQGLSSPVTKSFNRGEC-    217
7A9_WT         SIFPPSSEQLTSGGASVVCFLNNFYPKDINVKWKIDGSERQNGVLNSWTDQDSKDSTYSMSSTLTLTKDEYERHNSYTCEATHKTSTSPIVKSFNRNEC-    215
7A9_4DS        SIFPPSSEQLTSGGASVVCFLNNFYPKDINVKWKIDGSERQNGVLNSWTCQDSKDCTYSMSSTLTLTKDEYERHNSYTCEATHKTSTSPIVKSFNRNEC-    215
RTX_WT         FIFPPSDEQLKSGTASVVCLLNNFYPREAKVQWKVDNALQSGNSQESVTEQDSKDSTYSLSSTLTLSKADYEKHKVYACEVTHQGLSSPVTKSFNRGEC-    213
RTX_4DS        FIFPPSDEQLKSGTASVVCLLNNFYPREAKVQWKVDNALQSGNSQESVTCQDSKDCTYSLSSTLTLSKADYEKHKVYACEVTHQGLSSPVTKSFNRGEC-    213
5A12_WT        FIFPPSDEQLKSGTASVVCLLNNFYPREAKVQWKVDNALQSGNSQESVTEQDSKDSTYSLSSTLTLSKADYEKHKVYACEVTHQGLSSPVTKSFNRGEC-    215
5A12_6DS       CIFPPSDECLKSGTASVVCLLNNFYPREAKVQWKVDNALQSGNSQESVTCQDSKDCTYSLSSTLTLSKADYEKHKVYACEVTHQGLSSPVTKSFNRGEC-    215
2H11_WT        TLFPPSSEELQANKATLVCLISDFYPGAVTVAWKADSSPVKAGVETTTPSKQSN-NKYAASSYLSLTPEQWKSHRSYSCQVTHEGST--VEKTVAPTECS    215
2H11_4DS       TLFPPSSEELQANKATLVCLISDFYPGAVTVAWKADSSPVKAGVETTTPSCQSNCNKYAASSYLSLTPEQWKSHRSYSCQVTHEGST--VEKTVAPTECS    216
                :****.* * :. *::**::.:***   .* ** *.:  . .   :    :*:  .*: ** *:*:  ::: *. *:*:.**:  :  : *:.   **
```

**B**

```
E104v1_WT      EVQLVESGPGLVKPSETLSLTCTVSRFSLIG-YAITWIRQPPGKGLEWIGGISSAA-TTFYSSWAKSRVTISVDTSKNQFSLKLSSVTAADTAVYYCARDPRGYGAALDRLDLWGQGTLV    118
E104v1_2DS     EVQLVESGPGCVKPSETLSLTCTVSRFSLIG-YAITWIRQPPGKGLEWIGGISSAA-TTFYSSWAKSRVTISVDTSKNQFSLKLSSVTAADTAVYYCARDPRGYGAALDRLDLWGQGTLV    118
E104v1_4DS     EVQLVESGPGCVKPSETLSLTCTVSRFSLIG-YAITWIRQPPGKGLEWIGGISSAA-TTFYSSWAKSRVTISVDTSKNQFSLKLSSVTAADTAVYYCARDPRGYGAALDRLDLWGQGTCV    118
E104v1_6DS     EVQLVESGPGCVKPSETLSLTCTVSRFSLIG-YAITWIRQPPGKGLEWIGGISSAA-TTFYSSWAKSRVTISVDTSKNQFSLKLSSVTAADTAVYYCARDPRGYGAALDRLDLWGQGTCV    118
7A9_WT         EVQLVESGGGLVKPGGSLKLSCAASGFTFS-NYAMSWVRQTPEKRLEWVATISNGGRYTYYPDSVKGRFTISRDNAKNSLYLQMSSLRSEDTAMYYCARHLYRYD-VGGALDYWGQGTSV    118
7A9_4DS        EVQLVESGGGCVKPGGSLKLSCAASGFTFS-NYAMSWVRQTPEKRLEWVATISNGGRYTYYPDSVKGRFTISRDNAKNSLYLQMSSLRSEDTAMYYCARHLYRYD-VGGALDYWGQGTCV    118
RTX_WT         QVQLQQPGAELVKPGASVKMSCKASGYTFT-SYNMHWVKQTPGRGLEWIGAIYPGNGDTSYNQKFKGKATLTADKSSSTAYMQLSSLTSEDSAVYYCARST--YYGGDWYFNVWGAGTTV    117
RTX_4DS        QVQLQQPGAECVKPGASVKMSCKASGYTFT-SYNMHWVKQTPGRGLEWIGAIYPGNGDTSYNQKFKGKATLTADKSSSTAYMQLSSLTSEDSAVYYCARST--YYGGDWYFNVWGAGTCV    117
5A12_WT        EVQLVESGGGLVQPGGSLRLSCAASGFTIS-DYWIHWVRQAPGKGLEWVAGITPAGGYTYYADSVKGRFTISADTSKNTAYLQMNSLRAEDTAVYYCARFVFF---LPYAMDYWGQGTLV    116
5A12_6DS       EVQLVESGGGCVQPGGSLRLSCAASGFTIS-DYWIHWVRQAPGKGLEWVAGITPAGGYTYYADSVKGRFTISADTSKNTAYLQMNSLRAEDTAVYYCARFVFF---LPYAMDYWGQGTCV    116
2H11_WT        EVQLQESGPGLVKPPGTLSLTCAVSGGSISSSNWWSWVRQPPGKGLEWIGEIYHSG-STNYNPSLKSRVTISVDKSKNQFSLKLSSVTAADTAVYYCARGSSSWY-DLGPFDYWGQGTLV    118
2H11_4DS       EVQLQESGPGCVKPPGTLSLTCAVSGGSISSSNWWSWVRQPPGKGLEWIGEIYHSG-STNYNPSLKSRVTISVDKSKNQFSLKLSSVTAADTAVYYCARGSSSWY-DLGPFDYWGQGTCV    118
               :*** : *   *:*  :: ::* .*  ::       *::* * : ***:. *  .   * *    *.: *:: *.:..  :::.*: : *:*:*****          :: ** ** *

E104v1_WT      TVSSASTKGPSVFPLAPSSKSTSGGTAALGCLVKDYFPE-PVTVSWNSGALTSGVHTFPAVLQSSGLYSLSSVVTVPSSSLGTQTYICNVNHKPSNTKVDKKVEPKSCDKTHT-------    230
E104v1_2DS     TVSSASTKGPSVFPLAPSSKSTSGGTAALGCLVKDYFCE-PVTVSWNSGALTSGVHTFPAVLQSSGLYSLSSVVTVPSSSLGTQTYICNVNHKPSNTKVDKKVEPKSCDKTHTHHHHHHP    237
E104v1_4DS     TVSSASTKGPSVFPLAPSSKSTSGGTAALGCLVKDYFCECPVTVSWNSGALTSGVHTFPAVLQSSGLYSLSSVVTVPSSSLGTQTYICNVNHKPSNTKVDKKVEPKSCDKTHTHHHHHHP    238
E104v1_6DS     TVSSASTKGPSVCPLAPSSKSTSGGTACLGCLVKDYFCECPVTVSWNSGALTSGVHTFPAVLQSSGLYSLSSVVTVPSSSLGTQTYICNVNHKPSNTKVDKKVEPKSCDKTHTHHHHHHP    238
7A9_WT         TVSSAKTTAPSVYPLAPVCGDTTGSSVTLGCLVKGYFPE-PVTLTWNSGSLSSGVHTFPAVLQS-DLYTLSSSVTVTSSTWPSQSITCNVAHPASSTKVDKKIEPRGPTIK-P-------    228
7A9_4DS        TVSSAKTTAPSVYPLAPVCGDTTGSSVTLGCLVKGYFCECPVTLTWNSGSLSSGVHTFPAVLQS-DLYTLSSSVTVTSSTWPSQSITCNVAHPASSTKVDKKIEPRGPTIK-PHHHHHHP    236
RTX_WT         TVSAASTKGPSVFPLAPSSKSTSGGTAALGCLVKDYFPE-PVTVSWNSGALTSGVHTFPAVLQSSGLYSLSSVVTVPSSSLGTQTYICNVNHKPSNTKVDKKVEPKSCDKTHT-------    229
RTX_4DS        TVSAASTKGPSVFPLAPSSKSTSGGTAALGCLVKDYFCECPVTVSWNSGALTSGVHTFPAVLQSSGLYSLSSVVTVPSSSLGTQTYICNVNHKPSNTKVDKKVEPKSCDKTHTHHHHHHP    237
5A12_WT        TVSSASTKGPSVFPLAPSSKSTSGGTAALGCLVKDYFPE-PVTVSWNSGALTSGVHTFPAVLQSSGLYSLSSVVTVPSSSLGTQTYICNVNHKPSNTKVDKKVEPKSCDKTHT-------    228
5A12_6DS       TVSSASTKGPSVCPLAPSSKSTSGGTACLGCLVKDYFCECPVTVSWNSGALTSGVHTFPAVLQSSGLYSLSSVVTVPSSSLGTQTYICNVNHKPSNTKVDKKVEPKSCDKTHTHHHHHHP    236
2H11_WT        TVSSASTKGPSVFPLAPSSKSTSGGTAALGCLVKDYFPE-PVTVSWNSGALTSGVHTFPAVLQSSGLYSLSSVVTVPSSSLGTQTYICNVNHKPSNTKVDKKVEPKSCD----------    226
2H11_4DS       TVSSASTKGPSVFPLAPSSKSTSGGTAALGCLVKDYFCECPVTVSWNSGALTSGVHTFPAVLQSSGLYSLSSVVTVPSSSLGTQTYICNVNHKPSNTKVDKKVEPKSCDKTHTHHHHHHP    238
               ***:*.*..*** **** . .*:*.:. ******.** * ***::****:*:************ .**:*** *** **:  :*:  *** *  *.******:**:.
```

**C**

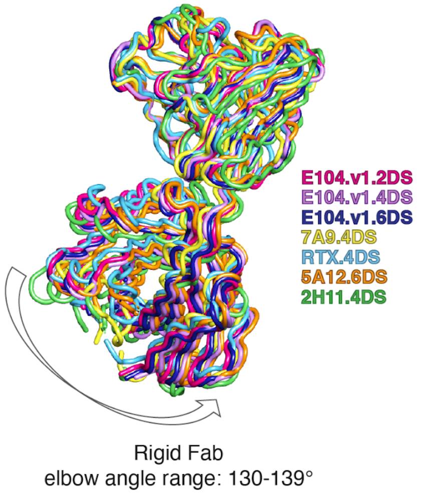

**Extended Data Fig. 9: Rigid Fab sequence and structural alignments. A and B.** Sequence alignments for the wild type (WT) and Rigid Fab variants of the light chains (A) and heavy chains (B) for the anti-tryptase Fab E104.v1, anti-Nav1.7 Fab 7A9, anti-CD20 Fab Rituximab

(RTX), anti-Ang2 Fab 5A12, and anti-KRAS Fab 2H11. Cysteine mutations introduced in the Rigid Fab 2DS, 4DS, and 6DS designs are highlighted in light blue, dark blue, and pink, respectively. **C.** VH-VL-based alignment illustrating the narrow range of elbow angles adopted by Rigid Fabs in the cryoEM structures from this study. Given the variety of fabs utilized in this study, it is likely that the elbow angles for each type of Fab, even with the same 4DS or 6DS designs are also determined by the type of the Fab (fully human or mouse, chimera, kappa vs lambda LC etc). Elbow angles were calculated using http://linum.proteinmodel.org/AS2TS/RBOW/index.html[27].

**Extended Data Table 1. Elbow angles of published rabbit Fab structures in the PDB.**

| PDB ID | LC | HC | Elbow angle (°) | Method | LC elbow disulfide |
|---|---|---|---|---|---|
| 4D3C | L | H | 137.6 | X-ray | no |
| 4HBC | L | H | 144 | X-ray | yes |
| 4HT1 | L | H | 137 | X-ray | no |
| 4JO1 | L | H | 142.7 | X-ray | yes |
| 4JO1 | M | I | 142.9 | X-ray | yes |
| 4JO2 | L | H | 140.8 | X-ray | yes |
| 4JO2 | M | I | 141.9 | X-ray | yes |
| 4JO3 | L | H | 154 | X-ray | yes |
| 4JO3 | M | I | 154.9 | X-ray | yes |
| 4JO4 | M | I | 153.5 | X-ray | yes |
| 4JO4 | L | H | 155.4 | X-ray | yes |
| 4MA3 | A | B | 139.8 | X-ray | no |
| 4MA3 | L | H | 142.6 | X-ray | no |
| 4O4Y | L | H | 144.8 | X-ray | yes |
| 4O51 | A | B | 148.9 | X-ray | no |
| 4O51 | C | D | 149.2 | X-ray | no |
| 4O51 | L | H | 149.5 | X-ray | no |
| 4O51 | E | F | 149.6 | X-ray | no |
| 4ZTO | M | I | 141.7 | X-ray | yes |
| 4ZTO | L | H | 143.6 | X-ray | yes |
| 4ZTP | L | H | 151.2 | X-ray | yes |
| 5C0N | D | C | 137.9 | X-ray | yes |
| 5C0N | L | H | 139.3 | X-ray | yes |
| 5DMG | D | C | 145.9 | X-ray | no |

| | | | | | |
|---|---|---|---|---|---|
| 5DMG | F | E | 151.3 | X-ray | no |
| 5DMG | L | H | 155 | X-ray | no |
| 5DRN | B | A | 150.3 | X-ray | yes |
| 5DRN | L | H | 150.8 | X-ray | yes |
| 5DS8 | L | H | 146.7 | X-ray | yes |
| 5DS8 | B | A | 156.4 | X-ray | yes |
| 5DSC | B | A | 148.7 | X-ray | yes |
| 5DSC | D | C | 148.8 | X-ray | yes |
| 5DSC | F | E | 149.7 | X-ray | yes |
| 5DSC | L | H | 151.5 | X-ray | yes |
| 5DTF | B | A | 153 | X-ray | yes |
| 5DTF | L | H | 154.4 | X-ray | yes |
| 5DUB | L | H | 146.7 | X-ray | yes |
| 5DUB | B | A | 156.5 | X-ray | yes |
| 5M63 | N | M | 143.3 | X-ray | yes |
| 5M63 | L | H | 143.4 | X-ray | yes |
| 5TH9 | L | H | 137.1 | X-ray | no |
| 5TH9 | M | I | 137.8 | X-ray | no |
| 5V6L | L | H | 139.2 | X-ray | yes |
| 5V6L | M | I | 149.1 | X-ray | yes |
| 5V6M | L | H | 140.5 | X-ray | no |
| 6CEZ | L | H | 138 | X-ray | yes |
| 6CJK | C | D | 146.1 | X-ray | yes |
| 6CJK | A | B | 149.4 | X-ray | yes |
| 6CWT | D | C | 140.7 | X-ray | no |
| 6CWT | B | A | 141.1 | X-ray | no |

| | | | | | |
|---|---|---|---|---|---|
| 6DID | H | L | 145.6 | EM | yes |
| 6I9I | B | A | 151.4 | X-ray | yes |
| 6I9I | L | H | 152.5 | X-ray | yes |
| 6LDV | L | H | 142.5 | X-ray | yes |
| 6LDW | A | B | 139.8 | X-ray | yes |
| 6LDW | L | H | 140.5 | X-ray | yes |
| 6LDX | L | H | 146.2 | X-ray | yes |
| 6LDY | L | H | 148.6 | X-ray | yes |
| 6LDY | B | A | 151.6 | X-ray | yes |
| 6T3F | L | H | 143.4 | X-ray | yes |
| 6X1S | F | E | 150.2 | X-ray | yes |
| 6X1S | D | C | 151.3 | X-ray | yes |
| 6X1S | L | H | 158.5 | X-ray | yes |
| 6X1S | B | A | 159.1 | X-ray | yes |
| 6X1T | D | C | 145.9 | X-ray | yes |
| 6X1T | K | J | 146.1 | X-ray | yes |
| 6X1T | B | A | 146.9 | X-ray | yes |
| 6X1T | L | H | 148.9 | X-ray | yes |
| 6X1T | F | E | 150.7 | X-ray | yes |
| 6X1T | I | G | 153.9 | X-ray | yes |
| 6X1U | L | H | 144.8 | X-ray | yes |
| 6X1U | B | A | 145.6 | X-ray | yes |
| 6X1V | L | H | 142.3 | X-ray | yes |
| 6X1W | L | H | 150.7 | X-ray | yes |
| 7D9Z | L | H | 144.7 | X-ray | yes |
| 7DCE | L | H | 144.7 | EM | yes |

| | | | | | |
|---|---|---|---|---|---|
| 7MFR | A | B | 148.7 | X-ray | no |
| 7MFR | X | Y | 148.7 | X-ray | no |
| 7NKS | D | C | 146.1 | X-ray | yes |
| 7NKS | L | H | 146.7 | X-ray | yes |
| 7NRH | L | H | 146.7 | EM | yes |
| 7O9S | L | H | 150.8 | X-ray | yes |
| 7RA7 | L | H | 147.3 | X-ray | yes |
| 7RA7 | B | A | 155.3 | X-ray | yes |
| 8DF1 | R | Q | 146.9 | X-ray | yes |
| 8DF1 | N | M | 147.7 | X-ray | yes |
| 8DF1 | V | U | 147.8 | X-ray | yes |
| 8DF1 | P | O | 148.3 | X-ray | yes |
| 8DF1 | L | H | 148.7 | X-ray | yes |

Elbow angles were calculated using http://linum.proteinmodel.org/AS2TS/RBOW/index.html [27]. LC and HC indicate chain IDs for the light and heavy chains, respectively.

**Extended Data Table 2. CryoEM data collection, refinement, and validation statistics for Tryptase-E104.v1 complexes.**

| | **Tryptase-E104.v1.WT** EMD-43200 PDB 8VGH | **Tryptase-E104.v1.2DS** EMD-43201 PDB 8VGI | **Tryptase-E104.v1.4DS** EMD-43202 PDB 8VGJ | **Tryptase-E104.v1.6DS** EMD-43203 PDB 8VGK |
|---|---|---|---|---|
| **Data collection and processing** | | | | |
| Magnification | 165,000 | 105,000 | 105000 | 105000 |
| Voltage (kV) | 300 | 300 | 300 | 300 |
| Detector | K2 | K3 | K3 | K3 |
| Electron exposure ($e^-$/Å) | 54.5 | 64.0 | 66.6 | 64.0 |
| Defocus range (μm) | 0.6-1.6 | 0.5-1.5 | 0.5-1.5 | 0.5-1.5 |
| Pixel size (micrographs) (Å) | 0.824 | 0.838 | 0.838 | 0.838 |
| Symmetry imposed | D2 | D2 | D2 | D2 |
| Initial particle images (no.) | 841,718 | 852,218 | 617,263 | 1,140,931 |
| Final particle images (no.) | 253,508 | 253,508 | 253,508 | 253,508 |
| Pixel size (map) (Å) | 1.0 | 1.0 | 1.0 | 1.0 |
| Map resolution (Å) | 2.9 | 2.7 | 2.5 | 2.4 |
| FSC threshold | 0.143 | 0.143 | 0.143 | 0.143 |
| Map resolution range (Å) | 2.9-18.8 | 2.7-21.4 | 2.5-19.4 | 2.4-19.7 |
| **Refinement** | | | | |
| Initial model used (PDB code) | 6VVU | 6VVU | 6VVU | 6VVU |
| Model resolution (Å) | 2.9 | 2.7 | 2.5 | 2.5 |
| FSC threshold | 0.5 | 0.5 | 0.5 | 0.5 |
| Model resolution range (Å) | 2.9-18.8 | 2.7-21.4 | 2.5-19.4 | 2.5-19.7 |
| Map sharpening B factor ($Å^2$) | -100 | -100 | -100 | -100 |
| Model composition | | | | |
| Non-hydrogen atoms | 20472 | 20480 | 20579 | 20567 |

| | | | | |
|---|---|---|---|---|
| Protein residues | 2668 | 2668 | 2672 | 2672 |
| Ligands | 0 | 0 | 0 | 0 |
| R.m.s. deviations | | | | |
| Bond lengths (Å) | 0.011 | 0.010 | 0.007 | 0.003 |
| Bond angles (°) | 0.944 | 0.884 | 0.826 | 0.768 |
| Validation | | | | |
| Molprobity score | 1.82 | 1.63 | 1.80 | 1.51 |
| Clashscore | 6.37 | 3.83 | 4.25 | 2.53 |
| Poor rotamers (%) | 0.13 | 0.70 | 1.57 | 0.30 |
| Ramachandran plot | | | | |
| Favored (%) | 92.41 | 92.75 | 93.11 | 92.27 |
| Allowed (%) | 7.02 | 6.98 | 6.59 | 7.54 |
| Disallowed (%) | 0.57 | 0.27 | 0.30 | 0.19 |
| Rama-Z score | | | | |
| Whole | -1.29 | -0.92 | -1.28 | -0.96 |
| Helix | -3.23 | -2.78 | -2.56 | -2.32 |
| Sheet | -0.13 | 0.10 | 0.22 | 0.54 |
| Loop | -1.01 | -0.86 | -1.39 | -1.26 |

**Extended Data Table 3. Data collection and refinement statistics for Rigid Fab crystal structures.**

| | **E104.v1.4DS.S112F** PDB 8VEG | **E104.v1.4DS.A114F** PDB 8VGE | **E104.v1.5DS** PDB 8VGF | **E104.v1.6DS** PDB 8VGG |
|---|---|---|---|---|
| **Data collection** | | | | |
| Space group | P 61 2 2 | P 61 2 2 | P 61 2 2 | P 61 2 2 |
| Cell dimensions | | | | |
| *a, b, c* (Å) | 116.34, 116.34, 136.05 | 115.10, 115.10, 137.83 | 115.52, 115.52, 136.42 | 115.49, 115.49, 136.04 |
| α, β, γ (°) | 90, 90, 120 | 90, 90, 120 | 90, 90, 120 | 90, 90, 120 |
| Resolution (Å) | 38.08 - 2.0 (2.072 - 2.0) | 46.869-2.010 (2.082 – 2.010) | 36.44 - 2.14 (2.216 - 2.14) | 36.42 – 2.71 (2.807-2.710) |
| $R_{sym}$ | 0.07047 (1.623) | 0.08241 (1.578) | 0.1421 (1.208) | 0.2333 (1.693) |
| *I/σI* | 27.52 (2.29) | 26.57 (2.27) | 15.14 (4.24) | 15.92 (2.42) |
| CC1/2 (%) | 1 (0.854) | 1 (0.808) | 0.998 (0.891) | 0.997 (0.799) |
| Completeness (%) | 99.05 (99.95) | 98.58 (99.64) | 99.93 (100.00) | 99.85 (100.00) |
| Redundancy | 19.3 (19.6) | 18.8 (19.5) | 15.6 (14.6) | 19.6 (19.7) |
| **Refinement** | | | | |
| Resolution (Å) | 2.0 | 2.01 | 2.14 | 2.71 |
| No. reflections | 717900 (71762) | 686885 (69671) | 471925 (43369) | 296265 (28711) |
| $R_{work}/R_{free}$ | 0.2147/0.2434 | 0.2037/0.2321 | 0.1791/0.2189 | 0.2066/0.2579 |
| No. atoms | 3428 | 3487 | 3468 | 3172 |
| Protein | 3228 | 3279 | 3225 | 3161 |
| Ligand/ion | N/A | N/A | N/A | N/A |
| Water | 200 | 208 | 243 | 11 |
| *B*-factors | 53.83 | 54.70 | 55.74 | 64.94 |
| Protein | 53.90 | 54.88 | 55.81 | 65.02 |
| Ligand/ion | N/A | N/A | N/A | N/A |
| Water | 52.70 | 51.85 | 54.89 | 41.51 |
| R.m.s. deviations | | | | |
| Bond length (Å) | 0.006 | 0.004 | 0.011 | 0.002 |
| Bond angles (°) | 1.25 | 0.69 | 1.28 | 0.60 |

**Extended Data Table 4. CryoEM data collection, refinement, and validation statistics for Nav1.7-7A9 and CD20-RTX complexes.**

| | **Nav1.7-7A9.WT**<br>EMD-43204<br>PDB 8VGL | **Nav1.7-7A9.4DS**<br>EMD-43208<br>PDB 8VGM | **CD20-RTX.WT**<br>EMD-43212<br>PDB 8VGN | **CD20-RTX.4DS**<br>EMD-43216<br>PDB 8VGO |
|---|---|---|---|---|
| **Data collection and processing** | | | | |
| Magnification | 165,000 | 165,000 | 165,000 | 165,000 |
| Voltage (kV) | 300 | 300 | 300 | 300 |
| Detector | Falcon 4 | Falcon 4 | Falcon 4 | Falcon 4 |
| Electron exposure ($e^-$/Å) | 44.0 | 44.0 | 44.0 | 44.0 |
| Defocus range (μm) | 0.8-1.8 | 0.8-1.8 | 0.8-1.8 | 0.8-1.8 |
| Pixel size (micrographs) (Å) | 0.731 | 0.731 | 0.731 | 0.731 |
| Symmetry imposed | C2 | C2 | C2 | C2 |
| Initial particle images (no.) | 1,752,500 | 1,312,570 | 1,262,843 | 1,580,660 |
| Final particle images (no.) | 391,335 | 362,778 | 390,855 | 375,469 |
| Pixel size (map) (Å) | 0.9357 | 0.9357 | 0.9357 | 0.9357 |
| Map resolution (Å) | 2.6 | 2.6 | 2.5 | 2.6 |
| FSC threshold | 0.143 | 0.143 | 0.143 | 0.143 |
| Map resolution range (Å) | 2.1-33.5 | 2.1-37.6 | 2.1-37.0 | 2.0-40.3 |
| **Refinement** | | | | |
| Initial model used (PDB code) | 6N4Q, 5EK0 | 6N4Q, 5EK0 | 6VJA | 6VJA |
| Model resolution (Å) | 2.6 | 2.6 | 2.6 | 2.6 |
| FSC threshold | 0.5 | 0.5 | 0.5 | 0.5 |
| Model resolution range (Å) | 2.1-33.5 | 2.1-37.6 | 2.1-37.0 | 2.0-40.3 |
| Map sharpening B factor ($Å^2$) | -100 | -100 | -100 | -100 |
| Model composition | | | | |
| Non-hydrogen atoms | 15990 | 15994 | 9150 | 9148 |
| Protein residues | 1960 | 1962 | 1204 | 1206 |
| Ligands | 0 | 0 | 0 | 0 |
| R.m.s. deviations | | | | |

| | | | | |
|---|---|---|---|---|
| Bond lengths (Å) | 0.012 | 0.003 | 0.012 | 0.005 |
| Bond angles (°) | 0.810 | 0.675 | 0.937 | 0.719 |
| Validation | | | | |
| Molprobity score | 1.66 | 1.43 | 1.59 | 1.41 |
| Clashscore | 5.27 | 3.13 | 4.28 | 4.37 |
| Poor rotamers (%) | 0.57 | 0.11 | 0.59 | 0.19 |
| Ramachandran plot | | | | |
| Favored (%) | 94.39 | 95.27 | 94.44 | 95.13 |
| Allowed (%) | 5.30 | 4.62 | 5.22 | 4.37 |
| Disallowed (%) | 0.31 | 0.10 | 0.34 | 0.50 |
| Rama-Z score | | | | |
| Whole | -1.45 | -1.36 | -0.64 | -0.21 |
| Helix | -0.89 | -1.06 | -0.93 | -1.08 |
| Sheet | -0.58 | -0.15 | 0.25 | 0.75 |
| Loop | -0.98 | -0.77 | -0.56 | -0.22 |

**Extended Data Table 5. CryoEM data collection, refinement, and validation statistics for Ang2 and KRAS complexes.**

| | **Ang2-5A12.6DS**<br>EMD-43220<br>PDB 8VGP | **KRAS-2H11.4DS**<br>EMD-43221<br>PDB 8VGQ |
|---|---|---|
| **Data collection and processing** | | |
| Magnification | 105,000 | 165,000 |
| Voltage (kV) | 300 | 300 |
| Detector | K3 | Falcon 4 |
| Electron exposure ($e^-$/Å) | 68.5 | 40.3 |
| Defocus range (µm) | 0.5-1.5 | 0.8-1.8 |
| Pixel size (micrographs) (Å) | 0.838 | 0.731 |
| Symmetry imposed | C1 | C1 |
| Initial particle images (no.) | 9,288,470 | 6,202,602 |
| Final particle images (no.) | 1,017,611 | 926,738 |
| Pixel size (map) (Å) | 1.0346 | 1.0573 |
| Map resolution (Å) | 2.7 | 2.8 |
| FSC threshold | 0.143 | 0.143 |
| Map resolution range (Å) | 2.3-41.0 | 2.4-39.9 |
| **Refinement** | | |
| Initial model used (PDB code) | 4ZFG | 7RP3 |
| Model resolution (Å) | 2.7 | 2.8 |
| FSC threshold | 0.5 | 0.5 |
| Model resolution range (Å) | 2.3-41.0 | 2.4-39.9 |
| Map sharpening B factor ($Å^2$) | -100 | -100 |
| Model composition | | |
| Non-hydrogen atoms | 5052 | 4672 |
| Protein residues | 648 | 600 |
| Ligands | 0 | MG (1), MKZ (1), GDP (1) |
| R.m.s. deviations | | |
| Bond lengths (Å) | 0.004 | 0.005 |
| Bond angles (°) | 0.703 | 0.849 |
| Validation | | |
| Molprobity score | 1.32 | 1.33 |
| Clashscore | 1.94 | 1.32 |
| Poor rotamers (%) | 0.91 | 2.14 |
| Ramachandran plot | | |
| Favored (%) | 94.69 | 96.62 |
| Allowed (%) | 5.16 | 3.04 |

| | | |
|---|---|---|
| Disallowed (%) | 0.16 | 0.34 |
| Rama-Z score | | |
| Whole | -0.77 | -0.31 |
| Helix | -2.85 | -1.97 |
| Sheet | 0.45 | 1.48 |
| Loop | -1.00 | -1.06 |

**Methods and Extended Data References**
